# A pathogenic AMPAR mutation disrupts hippocampal synapse maturation and homeostasis

**DOI:** 10.64898/2026.01.26.701845

**Authors:** Jian Xu, Yi-Zhi Wang, Toshihiro Nomura, Han M. Van, Yiwen Zhu, Charlotte C.M. Castillon, Morgane Chiesa, John J Marshall, John N Armstrong, Yongling Zhu, Jeffrey N. Savas, Anis Contractor

**Affiliations:** Department of Neuroscience, Northwestern University Feinberg School of Medicine, Chicago, IL 60613; Department of Neurology, Northwestern University Feinberg School of Medicine, Chicago, IL 60613; Department of Psychiatry and Behavioral Sciences, Northwestern University Feinberg School of Medicine, Chicago, IL 60613; Department of Neurobiology, Northwestern University Feinberg School of Medicine, Chicago, IL 60613

## Abstract

AMPA receptors (AMPARs) mediate fast excitatory synaptic transmission and are critical for activity-dependent synapse maturation and circuit development. *De novo* missense mutations in AMPAR subunits are linked to neurodevelopmental disorders (NDDs), yet how these variants disrupt neuronal function remains unclear. Here, we examine a recurrent pathogenic mutation in the GluA1 subunit (GRIA1 p.A636T) that alters AMPAR function and is associated with autism spectrum disorder (ASD) and intellectual disability (ID). Using GluA1^A636T^ knock-in mice of either sex, we identify impairments in hippocampal-dependent learning and working memory. Quantitative proteomics combined with bulk RNA-seq revealed age-dependent molecular remodeling in mutant hippocampus. Synaptic protein alterations were present in juvenile and adult mice, whereas a pronounced upregulation of mitochondrial and glial-associated proteins emerged more strongly in adulthood. Transcriptomic analyses showed partial concordance and highlighted divergence between RNA and protein regulation. Consistent with these molecular changes, there were increases in markers of oxidative stress in juvenile and adult mutant animals, whereas altered mitochondria morphology emerged selectively in adult mutants. Electrophysiological recordings demonstrated reduced basal excitatory transmission, persistence of silent synapses in adulthood and exaggerated activity-dependent synaptic potentiation, which were associated with reduced spine size consistent with impaired synapse maturation. Together, these findings indicate that a pathogenic AMPAR mutation disrupts developmental synapse maturation with secondary metabolic adaptations emerging over time.

**Significance Statement:** Mutations in genes encoding AMPA-type glutamate receptors are increasingly linked to neurodevelopmental disorders, including autism and intellectual disability. However, how these mutations disrupt brain function and development remains poorly understood. Using a mouse model carrying a recurrent disease-associated mutation in the GluA1 AMPA receptor subunit, we show that the presence of this mutant subunit protein impairs the normal maturation of excitatory synapses in the hippocampus. Mutant mice retain immature “silent” synapses into adulthood, exhibit abnormal synaptic plasticity, reduced spine size, and deficits in learning and memory. Molecular analyses reveal age-dependent remodeling of synaptic and mitochondrial proteins, suggesting that metabolic changes arise secondary to disrupted synapse development. These findings clarify how pathogenic AMPA receptor variants alter brain function over time.

## Introduction

AMPA receptors (α-amino-3-hydroxy-5-methyl-4-isoxazolepropionic acid receptors) mediate fast excitatory synaptic transmission making their assembly, trafficking, and regulation essential for supporting healthy brain function. (Shepherd and Huganir, 2007; Hansen et al., 2021). Through activity-dependent mechanisms, AMPARs also regulate synapse maturation and the development of neural circuits.

The importance of AMPAR function in human brain development is underscored by pathogenic *de novo* variants in all four of the genes encoding AMPAR subunits (*GRIA1–4*) which cause neurodevelopmental disorders (NDDs) including intellectual disability (ID) and autism spectrum disorder (ASD) (Geisheker et al., 2017; Salpietro et al., 2019; XiangWei et al., 2023). These conditions, collectively referred to as glutamate receptor ionotropic (GRI) disorders, frequently present with severe cognitive impairments. More broadly, *de novo* mutations in genes encoding synaptic proteins account for a substantial fraction of NDDs arising in simplex families (Iossifov et al., 2014). While the biophysical consequences of many disease-associated AMPAR variants have been characterized in heterologous systems, how altered receptor properties produce enduring disruptions of synapse development and neural function *in vivo* remains poorly understood.

Postnatal excitatory synapse maturation involves coordinated molecular and functional transitions. Newly formed synapses are initially “silent,” expressing NMDARs without functional AMPAR-mediated transmission, and subsequently undergo activity-dependent AMPAR recruitment (Isaac et al., 1995; Liao et al., 1995). Together with synaptic pruning these processes stabilize synaptic contacts and promote circuit maturation (Isaac et al., 1997; Stevens et al., 2007; Vardalaki et al., 2022). Disruptions in this developmental trajectory are implicated in multiple NDD models (Schwarz et al., 2026), yet how pathogenic alterations of AMPAR function affect synapse maturation remains unclear.

One recurrent pathogenic variant affects the GluA1 subunit at residue A636, within the conserved SYTANLAAF motif of the M3 transmembrane helix, commonly referred to as the “lurcher” site (Geisheker et al., 2017; Ismail et al., 2022; Tvergaard et al., 2024). Studies in recombinant systems have shown that substitutions at this site increase glutamate affinity, slow desensitization, and prolong channel deactivation kinetics, features typically interpreted as gain-of-function receptor gating (Klein and Howe, 2004). However, how these mutant receptors would affect complex neuronal properties including synapse maturation, circuit development, and neuronal homeostasis, in the intact brain remains unknown.

A further layer of complexity arises from the tight coupling between synaptic activity and neuronal bioenergetics (Li et al., 2004; Harris et al., 2012; Rangaraju et al., 2019). Although mitochondrial dysfunction is increasingly recognized as a common feature of NDDs (Rossignol and Frye, 2012; Liu et al., 2026), the temporal and mechanistic relationship between primary synaptic perturbations and subsequent mitochondrial alterations remains poorly defined (Anitha et al., 2023). In particular, it is unclear whether mitochondrial abnormalities represent an early driver of synapse dysfunction or a delayed consequence of altered circuit activity and homeostatic stress (Rojas-Charry et al., 2021; Anitha et al., 2023).

Here, we created a new knock-in mouse model expressing the pathogenic GluA1 A636T mutation to define how the presence of this mutation, *in situ*, reshapes synapse maturation and neuronal metabolic programs across development. By integrating MS-based quantitative proteomics and bulk transcriptomics with functional approaches, we uncover a staged molecular and functional response to mutant AMPAR expression. We show that the mutant GluA1 A636T subunit markedly reduces mature excitatory synapse properties, promotes the persistent retention of silent synapses into adulthood, alters synaptic plasticity in the hippocampus, remodels mitochondria morphology and elevates markers for oxidative and nitrosative stress. These synaptic-mitochondrion phenotypes are accompanied by impaired performance in spatial learning and working memory tasks.

Strikingly, discovery-based proteomic and bulk RNA sequencing analyses reveal that synaptic protein remodeling and signatures of aberrantly elevated metabolic activity emerge early in development, whereas robust alteration of mitochondrial proteins appear later and are not predicted by changes in the transcriptome. Together, these findings indicate that a pathogenic AMPAR mutation disrupts synapse maturation and is associated with delayed alterations in mitochondria, imposing lasting disruption on hippocampal function and providing new insight into the pathophysiology of GRI disorders.

## Methods

### Generation of GluA1^A636T^ mice

The CRISPR/Cas9 gene-editing system was used to generate the A636T amino-acid substitution in the mouse *Gria1* gene. The Alanine at position 636 (A636), encoded by exon 12, is specified by the codon GCC. In human patients, a G→A nucleotide substitution converts this codon to ACC, resulting in the A636T mutation. To introduce the equivalent mutation in mice, the endogenous GCC codon was changed to ACG, which also encodes threonine. These nucleotide substitutions additionally created a novel Tai I restriction enzyme site (ACGT^), enabling convenient genotyping (**Figure 1D-F**). Single-guide RNAs (sgRNAs), Cas9 mRNA, and a single-stranded oligodeoxynucleotide (ssODN) donor carrying the desired nucleotide substitutions were microinjected into mouse zygotes. Two founders, one female and one male, harboring the correctly edited A636T allele were identified and bred with wild-type C57BL/6J mice. Each founder produced viable heterozygous A636T mutant offspring (see **Figure 1E** for representative DNA sequencing chromatograms).

**Figure 1.**
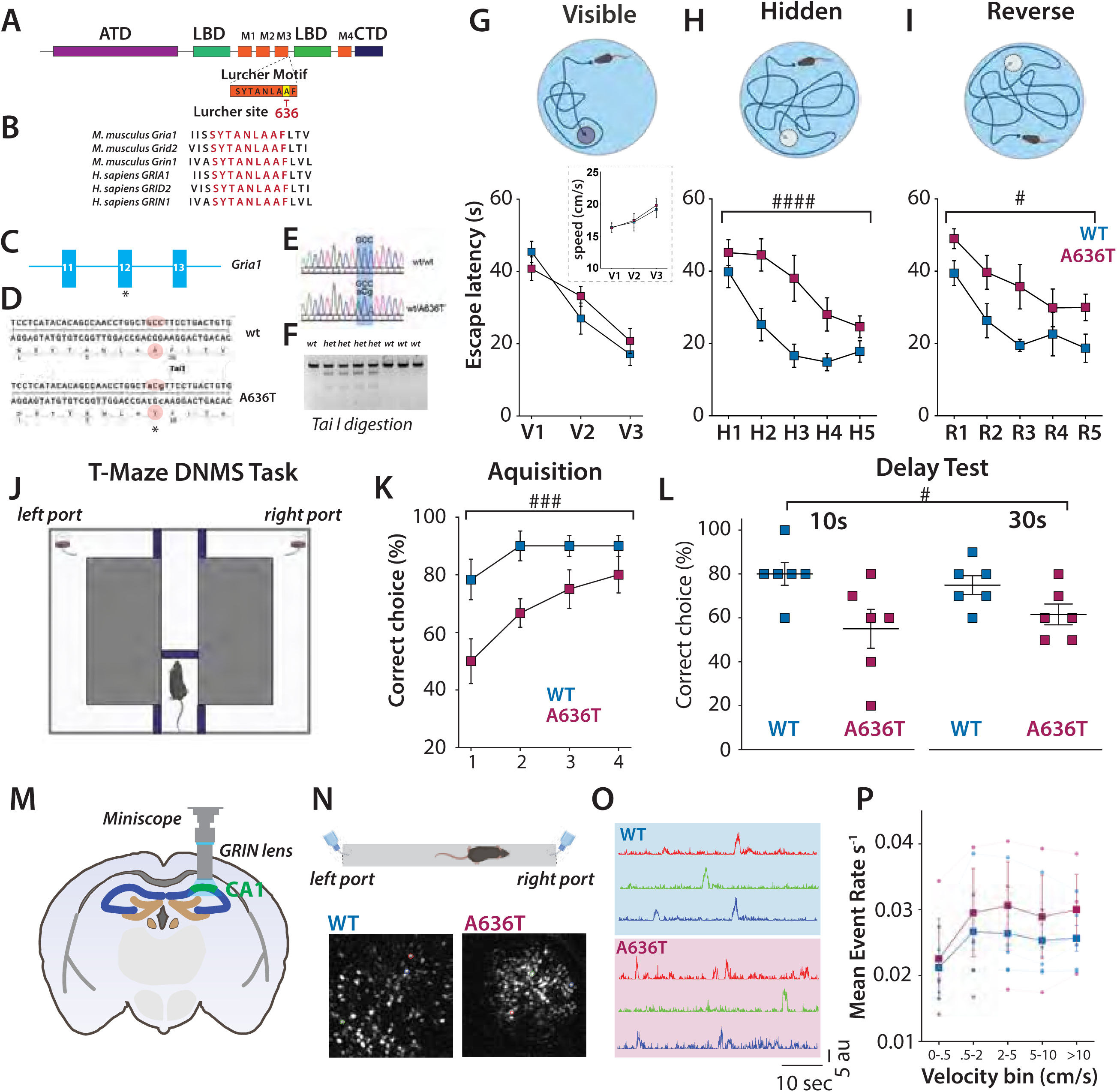
Generation and behavioral characterization of a GluA1^A636T^ knock-in mouse. **(A)** Linearized schematic of the GluA1 polypeptide highlighting the ligand-binding domain (LBD), transmembrane domains, and carboxyl-terminal domain (CTD). **(B)** Sequence alignment showing conservation of the SYTANLAAF (“lurcher”) motif across ionotropic glutamate receptors and species. **(C)** Schematic of the *Gria1* locus highlighting exon 12 and the location of the A636T mutation. **(D)** Sequence in exon 12 around the SYTANLAAF motif and site of editing **(E)** Sanger sequencing trace confirming heterozygous incorporation of the A636T mutation. **(F)** Introduction of a Tai I restriction site by the A636T mutation enables genotyping. **(G)** Escape latency during visible platform training in the Morris Water Maze. Inset, swimming speed during visible platform training. **(H)** Escape latency during hidden platform acquisition. **(I)** Escape latency during reversal learning. (WT, n = 10; GluA1^A636T^, n = 9). **(J)** Cartoon of the T-maze delayed non-match-to-sample maze. **(K)** Performance during training and acquisition with a 2-s delay. **(L)** Performance during working memory testing with a 10 second and 30 second delay. (WT, n = 6; GluA1^A636T^, n = 6). **(M)** Schematic of GRIN lens placement over dorsal CA1 for in vivo calcium imaging. **(N)** Linear track schematic and example field of view showing CA1 neurons from a WT and GluA1<u>^A636T^</u>mouse. **(O)** example Ca^2+^ traces during locomotion from cells extracted from fields shown in (N) for WT (top) and GluA1<u>^A636T^</u> (bottom). **(P)** Quantification of calcium event rates binned by velocity of the animals. ^####^P < 0.0001; ^###^P < 0.001, ^#^P < 0.05 (Two-Way Repeated Measures ANOVA). Data are represented as mean ± SEM. Error bars represent SEM.

### Behavioral Analysis

#### Morris Water Maze

Spatial learning and memory were assessed using the Morris Water Maze (MWM) as previously described (Xu et al., 2009; Xu et al., 2013). The apparatus consisted of a circular pool (120 cm diameter) filled with water maintained at 22–24°C and rendered opaque with non-toxic white paint. The pool was divided into four virtual quadrants, and distal visual cues were positioned around the testing room and remained constant throughout the experiment. A circular escape platform (10 cm diameter) was submerged 1 cm below the water surface and placed in a fixed quadrant during hidden platform training. Mouse behavior was recorded and analyzed using an automated video tracking system (LimeLight)

A total of 19 mice at ages of 10-12 weeks were tested (GluA1^A636T^ mice: n = 9; 4 females, 5 males; littermate controls: n = 10; 6 males, 4 females). Mice were first trained on a visible platform task to assess visual acuity, motivation, and motor function. The platform was marked with a visible cue and placed above the water surface. Mice received three trials per day for three consecutive days, starting from pseudo-randomized start locations. Each trial lasted a maximum of 60 seconds. Mice that failed to locate the platform within the allotted time were guided to it and allowed to remain on the platform for 15 seconds.

Following visible platform training, mice underwent hidden platform training for spatial learning. The platform was submerged and remained in a set location across training days. Mice received four trials per day for five consecutive days, with inter-trial intervals of at least 30 minutes. Escape latency and swim path length were recorded for each trial.

To assess cognitive flexibility and spatial memory updating, a reversal learning phase was performed in which the platform was relocated to the opposite quadrant. Mice received four trials per day for three consecutive days, and escape latencies were recorded as before during hidden platform training. Performance was quantified as escape latency to reach the platform. Swim speed was monitored to control for differences in motor performance and was not different across the performance of the task (**Figure 1G**).

#### T-maze delayed non-match-to-sample (DNMS) task

Spatial working memory was assessed using an automated T-maze delayed non-match-to-sample (DNMS) task (Moya et al., 2023; Webb et al., 2026). The maze consisted of a central start arm and two identical goal arms arranged in a T configuration. Automated sliding doors controlled access to each arm, and reward ports were located at the distal end of each goal arm (**Figure 1J**). Maze operation, trial structure, and data acquisition were controlled by custom software.

A total of 12 mice at ages of 10-12 weeks were tested (GluA1^A636T^ mice: n = 6; 3 females, 3 males; littermate controls: n = 6; 3 males, 3 females). Mice were habituated to the maze environment and trained to retrieve chocolate pellets from the goal arms prior to DNMS testing. Animals were food-restricted for 24 hours before training sessions to ensure motivation, while body weight was maintained above 85% of baseline. Habituation continued until mice reliably traversed the maze and consumed rewards without hesitation. Each DNMS trial consisted of a sample phase followed by a choice phase, separated by a delay period. During the sample phase, one goal arm was blocked and mice were allowed to enter the open arm to obtain a chocolate pellet reward. Following reward consumption, mice were confined to the start arm for the duration of the delay. During the subsequent choice phase, both goal arms were opened, and mice were rewarded for entering the arm opposite to the one visited during the sample phase (non-match rule). The identity of the sample arm was pseudorandomized across trials to minimize the development of side bias. Mice were trained on the DNMS task over four consecutive days using a 2-second delay between the sample and choice phases. Each training session consisted of 10 sets of a forced trial paired with a subsequent choice trial. Task acquisition was quantified as the percentage of correct non-match choices per session.

Following completion of training, working memory performance was assessed under increased cognitive demand by extending the delay period to 10 and 30 seconds. Performance during the extended delay condition was quantified as the percentage of correct choices and compared across genotypes.

### Histochemistry

#### General methods

Mice were decapitated, and brains were rapidly removed from the skull and post-fixed overnight at 4 °C in PBS containing 4% paraformaldehyde (PFA). Coronal brain sections (50 µm thick) were prepared using a Leica vibratome. Sections were collected and blocked for 60 min at room temperature in PBS containing 5% normal donkey serum (NDS; Jackson ImmunoResearch Labs, 017-000-121) and 1% Triton X-100. Sections were then incubated overnight at 4 °C with primary antibodies diluted in blocking solution. After primary antibody incubation, sections were washed three times for 15 min each in PBS and incubated with Alexa Fluor–conjugated secondary antibodies for 1 h at room temperature. Sections were washed three times and mounted using ProLong Gold Antifade Mountant with DAPI (Thermo Fisher Scientific). Images were acquired using a Zeiss epifluorescence microscope or a Nikon confocal microscope.

#### 3-Nitrotyrosine (NT) immunohistochemistry

Nitrotyrosine, a marker of nitrosative stress and inflammation, was detected using immunohistochemistry procedures described above. The primary antibody used was mouse anti-nitrotyrosine (1:1000; Abcam, ab61392).

#### Dihydroethidium (DHE) histochemistry

In situ visualization of superoxide (O₂⁻) and O₂⁻-derived oxidant production was assessed using dihydroethidium (DHE) histochemistry. Mice were injected intraperitoneally with 200 µl of PBS containing DHE (1 µg/µl; Thermo Fisher Scientific, cat. #D11347) with 1% DMSO as the vehicle control. Brains were harvested 30 min after injection, post-fixed overnight at 4 °C in PBS containing 4% paraformaldehyde (PFA), and then coronal sections (50 µm thick) were prepared using a vibratome. Oxidation of DHE, indicated by ethidium accumulation, was visualized by fluorescence microscopy (excitation 510 nm; emission 580 nm).

### In vivo GCaMP Recordings

Male mice aged approximately 3 months were anesthetized under continuous 1.5-2% isoflurane and underwent surgical implantation of a gradient refractive index (GRIN) lens (1 mm diameter, 4 mm length) above the dorsal hippocampal CA1. The lens was precoated with 1 μL of a 1:1 aqueous mixture of silk fibroin (5% w/v) and AAV9-CamKIIa-jGCaMP8m-WPRE (titer ≥ 1×10¹³ vg/mL). The bottom center of the lens was implanted at the following coordinates relative to bregma: -2.1 mm anteroposterior, +2.1 mm mediolateral, and -1.35 mm dorsoventral. The lens was secured to the skull using opaque dental cement and protected with parafilm and removable adhesive caulk. Animals recovered under standard postoperative care for 3–5 weeks prior to baseplate attachment and mounting of the miniature microscope (UCLA V4) for CA1 recordings.

Before Ca^2+^ imaging, mice were handled for 5 minutes daily for three consecutive days and trained to traverse a 175-cm-long, 6-cm-wide linear track in alternating directions. A 1-s water reward was delivered alternately at each end of the track, with distal visual landmarks present in the testing room. After reaching criterion performance (120 laps within 60 minutes), mice were further trained to run on the track while wearing the head-mounted miniscope (without recording) until the same criterion was achieved again. Ca^2+^ imaging commenced the following day.

During imaging sessions, mice were allowed to traverse the track up to 120 times over a maximum duration of 1 hour per day for 5-10 consecutive days. The position of the animal on the linear track was captured using a high-speed overhead camera, while GCaMP8m fluorescence was simultaneously recorded with the head-mounted miniscope during a 20-minute session each day. Throughout the experiment, mice were maintained under water restriction and kept at ≥ 80% of their initial body weight. We measured the Ca^2+^ event frequency as a function of the velocity of the mouse to determine whether CA1 neurons have the same activity levels in the GluA1^A636T^ mice. We used ezTrack software with a UBS webcam to track the velocity of the mice (Pennington et al., 2019).

To precisely determine the rate of onset of calcium events, we smoothed z-scored traces with a 3 sample rolling mean, required an above threshold peak to be 2 samples wide, and required an individual calcium event to decay to 1.4X the baseline value for 5 samples before another peak would be detected. We also defined the onset and offset of each calcium event as a 3 sample period 1.25X the baseline preceding and following the peak, respectively. To determine the instantaneous event rate, the number of onsets in a one second forward looking window (i.e. convolution of event counts per sample with a 1 second “boxcar” kernel) was used.

### RNA-seq

Two independent cohorts of mice were used for the RNA-seq experiments. The first cohort consisted of 10–12-week-old mice (3 WT and 4 GluA1^A636T^, mixed gender). The second cohort consisted of 4-week-old mice (4 WT and 3 GluA1^A636T^ mice mixed gender). Mouse hippocampi were rapidly dissected and immediately snap-frozen at −80 °C until RNA extraction. Total RNA was extracted using the RNeasy Plus Mini Kit (Qiagen, #74134). The stranded mRNA-seq was conducted in the Northwestern University NUSeq Core Facility. Briefly, total RNA samples were checked for quality using RINs generated from Agilent Bioanalyzer 2100. RNA quantity was determined with Qubit fluorometer. The Illumina Stranded mRNA Library Preparation Kit (llumina, Cat # 20040534) was used to prepare sequencing libraries from 500 ng of high-quality RNA samples (RIN>7). The Kit procedure was performed without modifications. This procedure includes mRNA purification and fragmentation, cDNA synthesis, 3’ end adenylation, Illumina adapter ligation, library PCR amplification and validation. The Illumina NovaSeq X Plus sequencer (first batch, 10-12-week-old mice) and the Illumina HiSeq 4000 sequencer (second batch, 4-week-old mice) were used to sequence the libraries with the production of single-end, 50 bp reads at the depth of 20-25 M reads per sample.

RNA-seq analysis was performed with OneStopRNAseq workflow (Li et al., 2020). Briefly, raw reads of individual FASTQ files was checked for quality control with FastQC and MultiQC (Ewels et al., 2016). Single-end reads were aligned to Mus musculus genome (mm39/GRCm39), with star RNA-seq aligner (Dobin et al., 2013). Aligned exon fragments with mapping quality higher than 20 were counted toward gene expression with featureCounts (Liao et al., 2014). Differential expression (DE) analysis was performed with DESeq2 (Pang et al., 1989). Significant DE genes were filtered with the criterion of false discovery rate < 0.05.

RNA-seq data have been deposited at GEO under the accession numbers GSE318397 and GSE318400.

### Proteomics

Two cohorts of mice were used for the proteomics experiments. The first cohort consisted of 10–12-week-old mice (7 WT and 8 GluA1^A636T^ mice, mixed gender). The second cohort consisted of 4-week-old mice (8 WT and 7 GluA1^A636T^ mice, mixed gender). Hippocampal tissue were dissected, snap frozen in liquid nitrogen and stored at −80 °C until use. Dissected hippocampal tissue was homogenized in RIPA lysis buffer (50 mM Tris, 150 mM NaCl, 0.1% SDS, 1 mM EDTA, 0.5% sodium deoxycholate, 1% Triton X-100, 1 x protease inhibitor cocktail (Thermo Fisher Scientific, Cat # 78443), 1 x phosphatase inhibitor (Thermo Fisher Scientific, Cat # 78420), pH 7.4) with an electronic homogenizer (Glas-Col, Cat # 099C-K54). Then excess 10% SDS solution was added into each sample to make the final SDS concentration to 1%. After sonication with a probe sonicator (Qsonica) for 3 × 1 min, hippocampal homogenates were solubilized at 4 °C for 1 h with rotation. Insoluble components were removed by centrifuging at 14,000 × g for 30 min. Methanol chloroform precipitation was used to clean and precipitate proteins.

Protein pellets were resuspended in 8 M urea (Thermo Fisher Scientific, Cat# 29700) prepared in 100 mM ammonium bicarbonate solution (Fluka, Cat# 09830). Dithiothreitol (DTT, DOT Scientific Inc, Cat# DSD11000) was applied to a final concentration of 5 mM. After incubation at RT for 20 min, iodoacetamide (IAA, Sigma-Aldrich, Cat# I1149) was added to a final concentration of 15 mM and incubated for 20 min at RT in the dark. Excess IAA was quenched with DTT for 15 min. Samples were diluted with 100 mM ammonium bicarbonate solution and digested for 3 h with Lys-C protease (1:100, Thermo Fisher Scientific, Cat# 90307_3668048707) at 37°C. Trypsin (1:100, Promega, Cat# V5280) was then added for overnight incubation at 37°C with intensive agitation. The next day, reaction was quenched by adding 1% trifluoroacetic acid (TFA, Fisher Scientific, Cat# O4902-100). The samples were desalted using Peptide Desalting Spin Columns (Thermo Fisher Scientific, Cat# 89852). All samples were vacuum centrifuged to dry.

TMT labeling was based on our previously reported methods (Wang et al., 2024b; Wang et al., 2024a). C18 column-desalted peptides were resuspended with 100 mM HEPES pH 8.5 and the concentrations were measured by micro BCA kit (Fisher Scientific, Cat# PI23235). For each sample, 100 μg of peptide labeled with 16plex-TMTpro reagent (0.4 mg, dissolved in 40 μL anhydrous acetonitrile, Thermo Fisher Scientific, Cat# A44520) and made at a final concentration of 30% (v/v) acetonitrile (ACN). Following incubation at RT for 2 h with agitation, hydroxylamine (to a final concentration of 0.3% (v/v)) was added to quench the reaction for 15 min. TMT-tagged samples were mixed at a 1:1:1:1:1:1:1:1:1:1:1:1:1:1:1:1 ratio. Combined sample was vacuum centrifuged to dryness, resuspended, and subjected to Peptide Desalting Spin Columns (Thermo Fisher Scientific, Cat# 89852). We used a high pH reverse-phase peptide fractionation kit (Thermo Fisher Scientific, Cat# 84868) to get eight fractions (preclean with 5.0%, then 10.0%, 12.5%, 15.0%, 17.5%, 20.0%, 22.5%, 25.0% and 50% of ACN in 0.1% triethylamine solution). The high pH peptide fractions were directly loaded into the autosampler for MS analysis without further desalting.

Three micrograms of each sample were loaded using an autosampler with a Thermo Vanquish Neo UHPLC system onto a PepMap Neo Trap Cartridge (Thermo Fisher Scientific, 174500; diameter, 300 µm; length, 5 mm; particle size, 5 µm; pore size, 100 Å; stationary phase, C18) coupled to a nanoViper analytical column (Thermo Fisher Scientific, 164570; diameter, 0.075 mm; length, 500 mm; particle size, 3 µm; pore size, 100 Å; stationary phase, C18) with a stainless-steel emitter tip assembled on the Nanospray Flex Ion Source with a spray voltage of 2,000 V. An Orbitrap Ascend (Thermo Fisher Scientific) was used to acquire all the MS spectral data. Buffer A contained 99.9% H2O and 0.1% formic acid, and buffer B contained 80.0% acetonitrile, 19.9% H_2_O with 0.1% formic acid. For each fraction (load 2 μg of peptide), the chromatographic run was for 4 h in total with the following profile: 0–2% for 7 min, 7% for 1 min, 7-10% for 5 min, 10-25% for 160 min, 25-33% for 40 min, 33-50% for 7 min, 50-95% for 14 min and 99% for 6 min.

We used a multiNotch MS3-based TMT method to analyze all the TMT samples (Ting et al., 2011; Wang et al., 2024b; Wang et al., 2024a). The scan sequence began with an MS1 spectrum (Orbitrap analysis, resolution 120,000, 400–1600 Th, AGC target 4×105, maximum injection time, auto). MS2 analysis, ‘Top speed’ (3 s), Collision-induced dissociation (CID, quadrupole ion trap analysis, AGC standard, NCE 35, maximum injection time 35 ms). MS3 analysis, top ten precursors, fragmented by HCD prior to Orbitrap analysis (NCE 55, max AGC 5×10^4^, maximum injection time 200 ms, isolation specificity 0.7 Th, resolution 45,000).

Protein identification/quantification and analysis were performed with Integrated Proteomics Pipeline - IP2 (Bruker, Madison, WI) using ProLuCID,66,67 DTASelect2,81,82 Census and Quantitative Analysis (Eng et al., 1994; Tabb et al., 2002; Xu et al., 2015). Spectrum raw files were extracted into MS1, MS2 and MS3 files using RawConverter (http://fields.scripps.edu/downloads.php). The tandem mass spectra were searched against UniProt mouse (downloaded on 07-25-2023) protein databases and matched to sequences using the ProLuCID/SEQUEST algorithm (ProLuCID version 3.1) with 5 ppm peptide mass tolerance for precursor ions and 600 ppm for fragment ions. The search space included all fully and half-tryptic peptide candidates within the mass tolerance window with no-miscleavage constraint, assembled, and filtered with DTASelect2 through IP2. To estimate peptide probabilities and false-discovery rates (FDR) accurately, we used a target/decoy database containing the reversed sequences of all the proteins appended to the target database. Each protein identified was required to have a minimum of one peptide of minimal length of six amino acid residues; however, this peptide had to be an excellent match with an FDR <1% and at least one excellent peptide match. After the peptide/spectrum matches were filtered, we estimated that the peptide FDRs were ≤1% for each sample analysis. We used Census and Quantitative Analysis in IP2 for protein quantification of TMT-MS experiments and protein quantification was determined by summing all TMT report ion counts. Proteins encoded by the same gene were collapsed into one entry, which were inferred by at least two given peptides quantified. Static modification: 57.02146 C for carbamidomethylation, 304.2071 for 16-plex TMTpro tagging; differential modifications: 15.9949 M for oxidation on M, 304.2071 for N-terminal 16-plex TMTpro tagging, 42.0106 for N-terminal Acetylation.

The mass spectrometry data presented in this study was deposited in Mass Spectrometry Interactive Virtual Enviroment (MassIVE) under the identifier MSV000100702 and ProteomeXchange under the identifier PXD074003.

### Plasmid construction, AAV packaging and systemic injection

AAVs were packaged, purified, and quantified as previously described (Xu et al., 2020; Xu et al., 2022). Briefly, AAVs were produced by polyethylenimine (PEI)–mediated transfection of HEK293 cells maintained in adherent culture using an AAV transfer plasmid, an AAV packaging plasmid (PHP.eB), and the adenoviral helper plasmid pAdΔF6. The AAV transfer plasmid, pAAV-hSyn-mito-EGFP, was constructed by cloning a 906-bp BglII (filled)–KpnI fragment containing the mito-EGFP sequence from pEF mito-cGFP (matrix) (Addgene #188904) into the KpnI–EcoRV backbone of pAAV-hSyn-EYFP (Addgene #117382). The final construct was verified by Sanger sequencing. The AAV packaging plasmid pUCmini-iCAP-PHP.eB was obtained from Addgene (#103005); this plasmid encodes the PHP.eB capsid, which enables efficient blood–brain barrier penetration and facilitates systemic AAV delivery to the central nervous system (Chan et al., 2017). At 96 h post-transfection, AAVs from both cell pellets and culture media were collected and purified by iodixanol gradient ultracentrifugation (Xu et al., 2022). Viral preparations were concentrated, formulated in PBS, and genome titers were determined using a nucleic acid dye– based method (Xu et al., 2020).

### CA1 mitochondria imaging

An AAV vector carrying the PHP.eB capsid (pAAV-hSyn-mito-EGFP) was delivered to GluA1^A636T^ mice and littermate control mice via the retro-orbital (RO) route (Chan et al., 2017). Approximately 1 µL of AAV (1.0 × 10¹² gc/mL) was diluted in 100 µL of PBS and injected into each mouse. Two age groups were examined. For the 4-week group (n = 8, 4 WT mice and 3 GluA1^A636T^ mice, mixed gender), RO injection was performed at postnatal day 20 (P20), and mice were sacrificed at P30. For the 12-week group (n = 8, 4 WT mice and 4 GluA1^A636T^ mice, mixed gender), RO injection was performed at 8–10 weeks of age, and mice were sacrificed 2– 4 weeks later. Brains were collected, and coronal brain sections were prepared for immunohistochemistry, to enhance the EGFP fluorescence signal. Brain sections were immunostained with chicken anti-GFP primary antibody (1:1000; Abcam, ab13970), followed by donkey anti-chicken CF488A secondary antibody (1:1000; Biotium, 20166).

Confocal images were subsequently acquired using a Nikon AXR confocal microscope (Nikon Instruments, Tokyo, Japan). Hippocampal CA1 pyramidal neurons were identified based fluorescence signals. Images were acquired using a 60X objective with numerical-aperture of 1.42, under standard acquisition conditions. CF488A was excited with a 488-nm laser, and emitted fluorescence was collected using the appropriate detector and emission filter settings. The pinhole was set to one Airy unit to provide optimal optical sectioning. For each neuron, a z-stack spanning the somatic region was acquired with step of 0.154 μm. Image acquisition was performed using Nikon NIS-Elements software. All images were acquired under identical microscope settings. For mitochondrial morphology analysis, images were analyzed using Fiji (ImageJ) with the Mitochondria Analyzer plugin (Chaudhry et al., 2020). Briefly, maximum-intensity projections were generated from Z-stack images, and the resulting 2D images were saved as 8-bit TIFF files. Mitochondrial morphology was then analyzed using the Mitochondria Analyzer plugin according to the published protocol (Chaudhry et al., 2020). The following parameters were quantified: total mitochondrial number, mitochondrial area, aspect ratio, and form factor. Specifically, aspect ratio (AR) is a measure of mitochondrial elongation and is calculated as the ratio of the major axis to the minor axis of each mitochondrion. An aspect ratio of 1 indicates a perfectly round mitochondrion, whereas higher values indicate increasingly elongated mitochondria. Form factor (FF) is a measure of mitochondrial shape complexity that incorporates both elongation and boundary irregularity. It is calculated as: FF = (Perimeter²) / (4π × Area). A higher form factor indicates mitochondria that are more elongated, branched, or have more complex boundaries, whereas a value close to 1 corresponds to smooth, circular mitochondria.

### CA1 spine imaging

An AAV vector carrying the PHP.eB capsid (pAAV-hSyn-YFP) was delivered to GluA1^A636T^ mice and littermate controls via RO injection (Chan et al., 2017). To achieve sparse infection of hippocampal neurons, approximately 1 µL of AAV (1.0 × 10¹² gc/mL) was diluted in 100 µL PBS and injected into each mouse. Two age groups were examined. For the 4-week group (n = 8, 4 WT mice and 4 GluA1^A636T^ mice, mixed gender), mice received RO injections at postnatal day 20 (P20) and were sacrificed at P30. For the 12-week group (n = 8, 4 WT mice and 4 GluA1^A636T^ mice, mixed gender), mice were injected at 8–10 weeks of age and sacrificed 2–4 weeks later. Brains were collected, and coronal brain sections were prepared for immunohistochemistry, to enhance the EYFP fluorescence signal as described above.

Confocal imaging was performed using a Nikon AXR laser-scanning confocal microscope (Nikon Instruments, Tokyo, Japan). Hippocampal CA1 pyramidal neurons were identified based on CF488A signals. Images were acquired using a 60X objective with numerical-aperture of 1.42, under standard acquisition conditions. CF488A was excited with a 488-nm laser, and emitted fluorescence was collected using the appropriate detector and emission filter settings. The pinhole was set to one Airy unit to provide optimal optical sectioning. Confocal z-stack images were acquired from second-order apical dendrites located 100–200 μm distal to the soma. For each dendritic segment of interest, a z-stack spanning the entire depth was collected with a z-step size of 0.154 μm. Image acquisition was performed using Nikon NIS-Elements software.

For dendritic spine density and morphology analysis, images were analyzed using Fiji (ImageJ). Our analysis was adapted from a published ImageJ-based method for quantifying dendritic spine density and morphology in hippocampal neurons (Lin et al., 2023), with the modification that spines were manually identified throughout the entire z-stack rather than from a two-dimensional maximum-intensity projection. Although analysis of the full z-stack is more labor-intensive, it provides more accurate identification and quantification of individual spines. For each animal, 5–10 second-order apical dendritic segments (20–30 μm in length), obtained from different CA1 pyramidal neurons, were analyzed. Well-resolved dendritic segments were examined manually throughout the z-stack, and individual spines were outlined using the “Freehand Selection tool”. To prevent the same spine from being counted more than once, each spine ROI was assigned to the z-slice in which the spine exhibited the largest cross-sectional area or highest fluorescence intensity. The "Associate ’Show All’ ROIs with slices" option in the ROI Manager was unchecked so that each ROI was displayed across all z-slices while remaining assigned to its original slice. After all spines within a dendritic segment had been manually outlined, Fiji (ImageJ) was used to measure spine area. In addition, the length of the dendritic segment was also measured, and spine density was calculated as the number of spines per micrometer of dendrite.

### Transmission electron microscopy (TEM)

Brains were collected from 10-12-week-old mice (3 WT and 3 GluA1^A636T^). Coronal brain sections (1 mm thick) were prepared using a vibratome. The CA1 region of the dorsal hippocampus was dissected and further trimmed into ∼1 mm³ tissue blocks. Samples were then fixed 4% paraformaldehyde and 0.5% glutaraldehyde in 0.2 M cacodylate buffer, followed by post-fixation in 1% osmium tetroxide. Tissues were then embedded in Epon resin (EMBed-812). Ultrathin sections were prepared at the Northwestern University Center for Advanced Microscopy and Nikon Imaging Center. Images were acquired by first performing low-magnification imaging to identify regions of interest, followed by high-magnification imaging. High-magnification images were collected using TIA software.

For each mouse, 3–5 electron micrographs were acquired from the stratum radiatum at a distance of approximately 100-200 μm from from the border of the CA1 pyramidal cell layer. Image analysis was performed using Fiji (ImageJ) software. Original TIFF images were analyzed as 8-bit grayscale images. Individual mitochondria were manually outlined using the “Freehand Selection tool”and added to the ROI Manager for analysis. Morphological and intensity measurements were obtained using the Measure function. For each mitochondrion, we measured area, perimeter, and mean gray value. Shape descriptors, including circularity and roundness, were also measured. Circularity ranges from 0 to 1, with a value of 1 representing a perfect circle and values approaching 0 indicating increasingly elongated or irregularly shaped objects. Roundness also ranges from 0 to 1, with a value of 1 representing a perfect circle and values approaching 0 indicating increasingly elongated structures. Unlike circularity, which is influenced by both object elongation and boundary irregularity, roundness primarily reflects object elongation and is less sensitive to irregularities along the object boundary.

### Electrophysiology

#### Acute hippocampal slice preparation

Acute hippocampal slices were prepared from male and female mice at the indicated ages. Animals were deeply anesthetized with isoflurane and transcardially perfused with ice-cold, oxygenated cutting solution containing (in mM): 87 NaCl, 75 sucrose, 25 glucose, 25 NaHCO_3_, 2.5 KCl, 1.25 NaH_2_PO_4_, 7 MgCl_2_, and 0.5 CaCl_2_, equilibrated with 95% O_2_ / 5% CO_2_. Brains were rapidly removed and 350 µm thick transverse hippocampal slices were cut using a vibrating microtome (Leica VT1200S).

Slices were transferred to a holding chamber, and the cutting solution was gradually replaced with the holding solution containing artificial cerebrospinal fluid (ACSF; in mM: 125 NaCl, 25 NaHCO_3_, 25 glucose, 2.5 KCl, 1.25 NaH_2_PO_4_, 1 CaCl_2_, and 2 MgCl_2_). The holding solution also contained 10 μM DL-APV and 100 μM kynurenate to reduce excitotoxicity, and slices were incubated at 34°C for 30 minutes before being maintained at room temperature until recordings. Slices were transferred to the recording chamber after incubation and were perfused with normal ACSF containing the following (in mM): 125 NaCl, 2.4 KCl, 1.2 NaH_2_PO_4_, 25 NaHCO_3_, 25 glucose, 2 CaCl_2_, and 1 MgCl_2_ at 30 °C during recordings. All solutions were continuously bubbled with 95% O_2_ / 5% CO_2_.

#### Whole-cell voltage-clamp recordings

Whole-cell recordings were obtained from visually identified CA1 pyramidal neurons using infrared differential interference contrast (IR-DIC) microscope (Zeiss) equipped with Dodt contrast (Luigs & Neumann). Recording pipettes (3–5 MΩ) were filled with internal solution containing (in mM): 135 CsMeSO_3_, 10 TEA-Cl, 0.2 EGTA, 10 HEPES, 1 MgCl_2_, 10 Na-phosphocreatine, 2 Mg-ATP, 0.3 Na-GTP, and 5 QX-314 (pH 7.3, 290 mOsm). For miniature EPSC (mEPSC) recordings, neurons were voltage-clamped at −70 mV in the presence of tetrodotoxin (TTX, 0.5 µM) to block action potentials and picrotoxin (50 µM) to block GABA_A_ receptor–mediated currents. mEPSCs were detected and analyzed using Eventer software. Event amplitude, frequency, and decay kinetics were quantified from stable recording periods of 5 minutes.

#### Silent synapse recordings using minimal stimulation

Minimal stimulation experiments were performed to assess the abundance of functionally silent synapses. A monopolar stimulating electrode was placed in the *stratum radiatum* to activate Schaffer collateral (SC) input axons. Stimulation intensity was adjusted to evoke synaptic responses with a substantial failure rate, consistent with activation of one or a small number of presynaptic fibers. EPSCs were recorded at holding potentials of −70 mV to isolate AMPAR-mediated currents and +40 mV to record mixed AMPAR/NMDAR responses. For each cell, at least 25 and up to ∼100 consecutive stimuli were delivered at each holding potential at 0.2 Hz. Silent synapse abundance was estimated from the ratio of failure rates at −70 mV and +40 mV using established methods.

#### Alternate method for estimating silent synapse abundance

Silent synapse abundance was additionally estimated using an approach based on analysis of early and late components of evoked EPSCs (EPSC_E_ and EPSC_L_) under a modified minimal stimulation configuration (Isaac et al., 1995). Schaffer collateral inputs were stimulated extracellularly, and CA1 pyramidal neurons were voltage-clamped at a holding potential of +40 mV to permit detection of both AMPAR-and NMDAR-mediated responses. Stimulation intensity was initially adjusted to a level at which synaptic responses exhibited a high failure rate (typically >90%) and was then incrementally increased. At each stimulation intensity, 25 sweeps were acquired, and failure rates of EPSC_E_ and EPSC_L_ were quantified.

A higher failure rate of EPSC_E_ relative to EPSC_L_ indicates the presence of NMDAR-only (silent) synapses, as EPSC_E_ and EPSC_L_ are predominantly mediated by AMPARs and NMDARs, respectively (Isaac et al., 1995). Plotting EPSC_L_ failure rate against EPSC_E_ failure rate enables quantitative estimation of silent synapse abundance. When synapses are mature and contain both receptor types, the relationship approximates the unity line (y = x), with slope near 1 and intercept near 0.

#### Extracellular recordings

Field excitatory postsynaptic potentials (fEPSPs) were recorded from the CA1 subregion of the hippocampus. Stimulating (∼1 MΩ) and recording (2-3 MΩ) glass microelectrodes were filled with ACSF and placed in the *stratum radiatum*. Input–output (I-O) curves were generated by varying stimulus intensity and measuring the initial slope of the fEPSPs. Paired-pulse ratio (PPR) was estimated by measuring the slopes of the first and second fEPSPs elicited by two stimuli delivered with several interstimulus intervals of 50 – 900 ms.

#### Activity-dependent synaptic unsilencing and plasticity

To assess activity-dependent synaptic unsilencing, CA1 pyramidal neurons were voltage-clamped at −70 mV and minimal stimulation was delivered to Schaffer collateral inputs at 0.2 Hz. After acquisition of a stable baseline for 5 min (60 sweeps), a brief high frequency stimulus train (100 Hz for 0.5 s) was applied to induce synaptic potentiation. EPSC failure rates were quantified during the baseline period and again 25–30 min following train stimulation.

Long-term potentiation (LTP) was examined using an extracellular field recording configuration. After establishing a stable baseline for at least 10 min, the same short high frequency stimulation protocol (100 Hz for 0.5 s) was delivered to induce LTP. Potentiation magnitude was quantified as the percentage change in the slope of the field excitatory postsynaptic potential (fEPSP) relative to baseline.

Electrophysiological signals were recorded using a Multiclamp 700B amplifier (Molecular Devices), low-pass filtered at 2 kHz, and digitized at 10–50 kHz. Data acquisition and analysis were performed using pClamp 11 software (Molecular Devices).

## Data analysis and statistics

All data are presented as mean ± SEM unless otherwise indicated. Statistical analyses were performed using GraphPad Prism, DAVID, GSEA v4.4.0, and custom scripts in Python and/or R. Normality was assessed using the Shapiro–Wilk test when sample size permitted. For comparisons between two independent groups, unpaired two-tailed Student’s t-tests or Welch’s t-tests were used for normally distributed data, and Mann–Whitney U tests were used for nonparametric data. For paired comparisons, paired t-tests or Wilcoxon signed-rank tests were applied as appropriate. Experiments involving multiple groups or repeated measurements were analyzed using one-or two-way ANOVA or mixed-effects models with appropriate post hoc corrections (Tukey, Sidak, or Holm–Sidak). For proteomic data, p-values were adjusted using the Benjamini-Hochberg (BH) method. For GSEA assay, limma empirical Bayes moderated t-statistic was used to rank -omic data. Categorical data were analyzed using Fisher’s exact test when applicable. Significance was defined as p < 0.05. Exact statistical tests, sample sizes, and p values are reported in the figure legends and Results.

## Results

### Generation and functional characterization of the pathogenic GluA1^A636T^ knock-in mouse

To define the *in vivo* impact of a recurrent, pathogenic AMPAR variant linked to neurodevelopmental disorders, we generated a *Gria1* knock-in mouse carrying the GluA1 A636T substitution at the endogenous locus (**Figure 1**). This variant affects a highly conserved residue within the SYTANLAAF motif of the M3 transmembrane helix, a critical region involved in AMPAR gating (**Figure 1A**). Heterozygous GluA1^A636T/+^ mice are viable and born at Mendelian ratios, whereas homozygous mutants were not recovered, consistent with a dominantly acting pathogenic allele and associated early lethality. Both DNA sequencing and the introduction of a silent Tai I site confirmed that the introduced mutation was present in the mice (**Figure 1C-F**).

Given the central role of GluA1-containing AMPARs in hippocampal synapse maturation and spatial memory, we first asked whether pathogenic GluA1A636T expression disrupts hippocampal-dependent behaviors in adult mice. In the Morris Water Maze, GluA1^A636T^ mice (n = 9; 4 females, 5 males, 10-12 weeks) displayed significantly prolonged escape latencies during acquisition of the hidden platform task compared to littermate controls (n = 10; 6 males, 4 females, 10-12 weeks) (Two-Way Repeated Measures ANOVA , F_1,17_ =26.23, P < 0.0001) (**Figure 1H**), despite intact performance during visible platform training (F_1,17_ = 0.5274, P = 0.4776), and comparable swimming speeds between genotypes (F₁,₉ = 0.055, P = 0.8186), indicating preserved sensorimotor function (**Figure 1G, insert**). Mutant mice also exhibited impaired performance during reversal learning (F_1,17_ = 7.271, P = 0.0153), suggesting reduced cognitive flexibility (**Figure 1I**).

To further assess hippocampal working memory, we tested the mice in an automated T-maze delayed non-match-to-sample (DNMS) task that requires the transient maintenance of spatial information across a variable delay (**Figure 1J**). Mice were first trained in the maze to choose the correct arm (alternate arm to previously visited arm) with an introduced delay to test working memory. During the 4-day training period with a 2-second delay, the control group (n = 6; 3 males, 3 females) consistently outperformed the mutant group (n = 6; 3 males, 3 females), even though the mutant mice exhibited improvement over the course of the training and eventually performed similarly (Two-Way Repeated Measures ANOVA, F_1,10_ = 12.81, P = 0.0050) (**Figure 1K**). After acquisition training, the delay period was extended to 10 seconds and then to 30 seconds to test the working memory component. Across both the 10 s and 30 s delay tests, GluA1^A636T^ mice performed significantly worse than WT littermates (Two-way repeated-measures ANOVA, main effect of genotype: F_1,10_ = 9.03, P = 0.0132). There was no effect of delay (F_1,10_ = 0.02, P = 0.886) and no genotype × delay interaction (F_1,10_ = 1.05, P = 0.329), indicating that the impairment was evident across both delay intervals (**Figure 1L**). Together, these behavioral phenotypes reveal robust deficits in hippocampal-dependent learning and working memory.

To determine whether the GluA1^A636T^ mutation alters overall hippocampal neuronal activity *in vivo*, we performed cellular Ca^2+^ imaging of CA1 pyramidal neurons using head-mounted miniscopes. Mice were implanted with a gradient refractive index (GRIN) lens over the dorsal hippocampus and expressed GCaMP8m under the CaMKIIα promoter, allowing longitudinal monitoring of neuronal activity during behavior (**Figure 1M**). Following recovery and habituation to the miniscope, mice traversed a linear track to obtain water rewards while position and Ca^2+^ signals were recorded simultaneously (**Figure 1N & O**). This paradigm was used to assess overall neuronal activity during a simple locomotor behavior. Ca^2+^ event rates varied significantly with locomotor velocity in both genotypes (**Figure 1P**), consistent with previous studies demonstrating modulation of CA1 neuronal activity by running speed (Dombeck et al., 2010). However, event rates did not differ significantly between WT and GluA1^A636T^ mice across velocity bins (Two-way repeated-measures ANOVA; main effect of velocity: F_4,28_ = 16.70, P < 0.0001; main effect of genotype: F_1,7_ = 0.40, P = 0.549; genotype × velocity interaction: F_4,28_ = 0.87, P = 0.492) (**Figure 1P**). These findings indicate that the pathogenic mutation does not produce a detectable large-scale alteration in overall CA1 calcium event frequency during simple locomotor behavior.

### Quantitative hippocampal proteomics and bulk RNAseq reveal robust, age-specific remodeling in juvenile and adult GluA1^A636T^ mice

To define how expression of the pathogenic GluA1A636T subunit reshapes the hippocampal proteome, we performed tandem mass tag (TMT)-MS-based quantitative proteomic analyses using mice at two ages (McAlister et al., 2014). Hippocampal tissue was collected from juvenile (4-week-old, n = 8 WT and n = 8 GluA1^A636T^, mixed gender) and adult (10-12 week-old, n = 7 WT and n = 8 GluA1^A636T^, mixed gender) mice (**Figure 2A**). t-Distributed stochastic neighbor embedding (tSNE) analysis demonstrated clear clustering of biological replicates by genotype at both ages, indicating reproducible proteomic differences between WT and GluA1^A636T^ hippocampus (**Figure 2B**). GluA1^A636T^ expression significantly altered the hippocampal proteomes of both juvenile and adult mice (**Datasets S1 & 2**). Notably, the impact on the adult hippocampal proteome was more robust. In total we quantified 6113 proteins in 4-week old datasets and 5380 proteins in 10-week old datasets. Differentially expressed proteins (DEPs) (BH-adjusted p-value < 0.05) diverged markedly between juvenile and adult samples (**Figure 2C**). While only ∼1.6% quantified proteins are DEPs in the 4-week-old dataset (**Figure 2C**, top panel), DEPs are ∼27.8% of total quantified proteins in the 10-week-old dataset (**Figure 2C**, lower panel). The adult dataset also showed more than eight-fold more upregulated DEPs (DEP-Up, n = 1340) than downregulated DEPs (DEP-Dn, n = 158).

**Figure 2.**
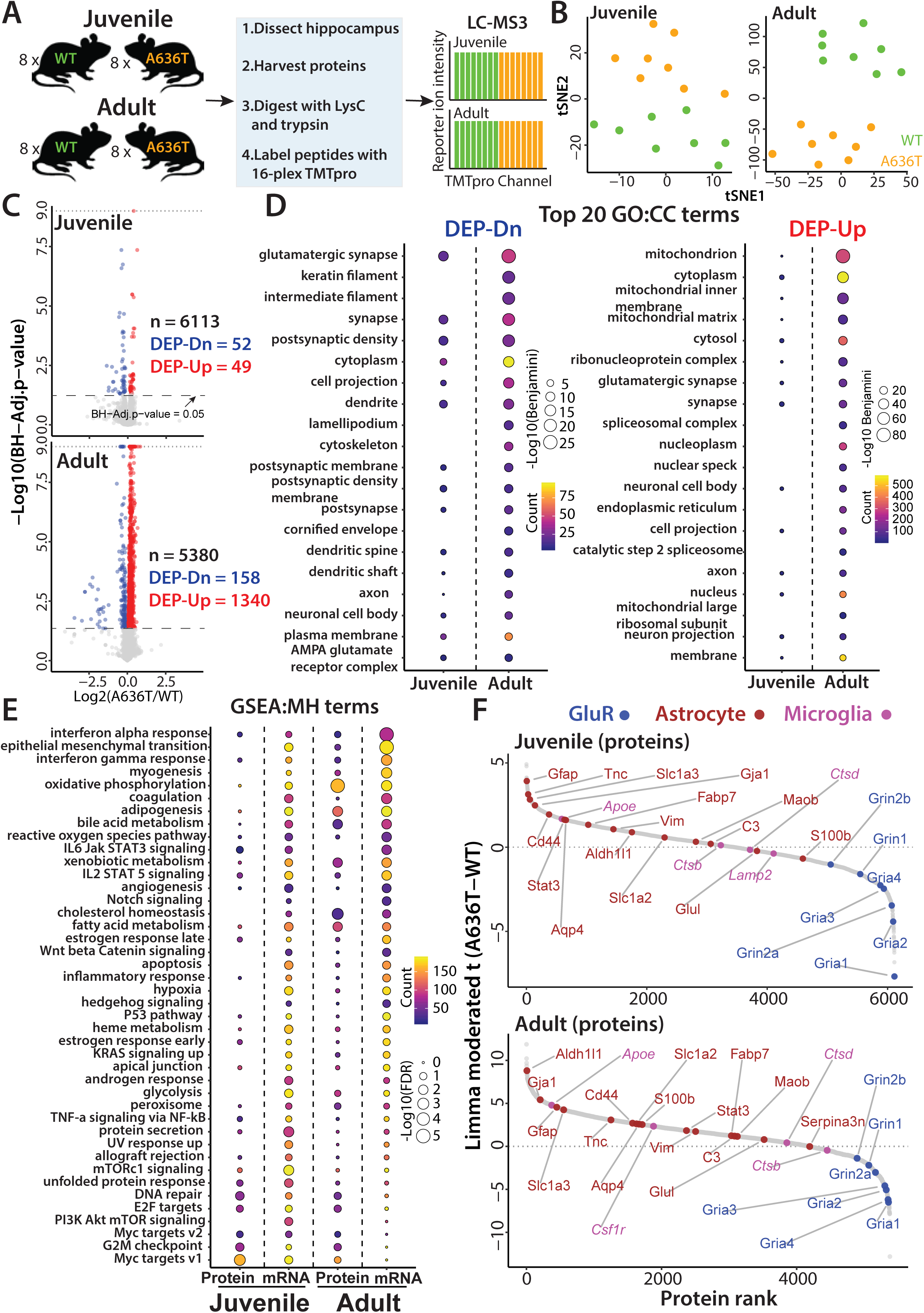
Developmental proteomic and transcriptomic profiling reveal age-dependent synapse and mitochondrial alterations in GluA1^A636T^ hippocampus. (**A**) Schematic of two independent TMT-MS proteomic experiments performed in juvenile and adult hippocampus. (**B**) t-Distributed stochastic neighbor embedding (tSNE) of TMT-MS profiles showing separation of biological replicates by genotype. Each group: WT, n = 8; GluA1^A636T^, n = 8. (**C**) Volcano plot of GluA1^A636T^-associated proteomic changes in juvenile (top panel) and adult mice (lower panel). Upregulated differentially expressed proteins (DEP-Up) are shown in red and downregulated proteins (DEP-Dn) in blue. Differential expression was defined as BH-adjusted p-value < 0.05. (**D**) Gene annotation analysis of DEP-Dn (left panel) and DEP-Up (right panel). Top 20 significantly enriched GO:CC terms in adult samples, shown alongside the corresponding terms in juvenile samples. (**E**) Gene set enrichment analysis (GSEA) comparing proteomics with bulk RNA-seq. Significantly enriched MSigDB Hallmark (MH) terms (FDR < 0.25 at least in one omic dataset) are shown alongside the same terms in proteomics and transcriptomics, consistent with increased ROS stress, inflammatory signaling, and apoptosis-related programs in adult hippocampus. (**F**) Limma moderated t-based ranking plots highlighting astrocyte activation signatures in both juvenile and adult GluA1^A636T^ hippocampus. See Datasets S1-S3 for detailed results of the proteomic and bulk RNA-sequencing analyses.

To interpret the biological events underlying these proteomic alterations, we performed a series of bioinformatic analyses. Gene Ontology (GO) cellular component (CC) enrichment of the DEPs (**Figure 2D, Datasets S1 & 2**) indicated that, in the juvenile hippocampus, the effects of GluA1^A636T^ expression are largely concentrated on synapses and dendrites, with most DEP-Dn corresponding to postsynaptic and dendritic protein elements (**Figure 2D**, left panel). In contrast, in the adult hippocampus, GluA1^A636T^ drove a markedly stronger reduction of postsynaptic and dendritic proteins and broadly decreased proteins across multiple synaptic compartments and cytoskeletal assemblies. Notably, these decreases were accompanied by pronounced elevated fold change of mitochondrial and nuclear proteins in the DEP-Up proteins (**Figure 2D**, right panel). Together, the GO:CC profiles suggest selective postsynaptic dysregulation in juveniles, whereas adults exhibit more global neuronal disturbance, accompanied by a broad elevation of protein levels and signatures of mitochondria emerging only in the adult samples.

To further investigate if the elevated protein levels observed in the GluA1^A636T^ mouse hippocampus can be explained by concordant gene transcription changes, we performed bulk RNA-seq on hippocampal tissue from 4-week-old (n = 3 WT and n = 4 GluA1^A636T^) and 10-week-old mice (n = 4 WT and n = 3 GluA1^A636T^) (**Figure 2E & Figure S1A & B**). We then conducted Gene Set Enrichment Analysis (GSEA) using MSigDB hallmark (H) gene sets, integrating the proteomic and RNA-seq datasets (**Figure 2E & S1C, Datasets S1-3**)(Mootha et al., 2003; Subramanian et al., 2005). Overall, the transcriptomic and proteomic profiles were highly concordant. Notably, the analyses indicated signatures of reactive oxidative stress (ROS), DNA repair and apoptosis related pathways already present at the juvenile stage (**Figure 2E**). In the adult mice, these signatures become more pronounced, consistent with an accumulation of cellular stress response in the hippocampus. This suggests that the GluA1^A636T^ variant imposes an early metabolic and proteostasis burden, and over time that chronic strain progresses into mitochondrial dysfunction and elevated ROS, ultimately driving secondary inflammatory remodeling in adult hippocampus. Finally, as astrocytes and microglia can both contribute to oxidative stress and neuroinflammation (Block et al., 2007; Chen et al., 2020; Giovannoni and Quintana, 2020; Patani et al., 2023), we applied a limma moderated t statistic-based ranking analysis to examine signature proteins for these glial populations (**Figure 2F**) (Smyth, 2004; Ritchie et al., 2015). There are a substantial number of reactive astrocyte markers and effectors found in this dataset including Gfap, Vim, Stat3, C3, Cd44, Serpina3n, Apoe, and Tnc, which were robustly increased in both juvenile and adult hippocampus consistent with an astrogliosis-associated cytoskeletal, inflammatory / complement, and ECM-remodeling program (**Figure 2F**). In contrast, microglia signature proteins showed comparatively modest and less consistent changes.

Collectively, these findings define a developmental pathological progression in GluA1^A636T^ mice in which early synaptic protein alterations are accompanied by increased oxidative stress signatures in the juvenile mice, as supported by both transcriptomic and proteomic analyses. As animals mature, these changes become more pronounced and are associated with the emergence of mitochondrial protein alterations, ROS accumulation, and broader molecular remodeling in the adult hippocampus. Together, these results indicate a sustained, evolving cellular stress response across development.

### Pathogenic GluA1^A636T^ expression is associated with progressive remodeling of mitochondrial morphology

Proteomic analysis of adult GluA1^A636T^ mouse hippocampus revealed increased abundance of numerous mitochondrial proteins (**Figure 3A, Dataset S4**) Because mitochondrial dysfunction has been implicated in both oxidative stress and inflammatory signaling, we next asked whether these molecular changes were accompanied by alterations in mitochondrial morphology.

**Figure 3.**
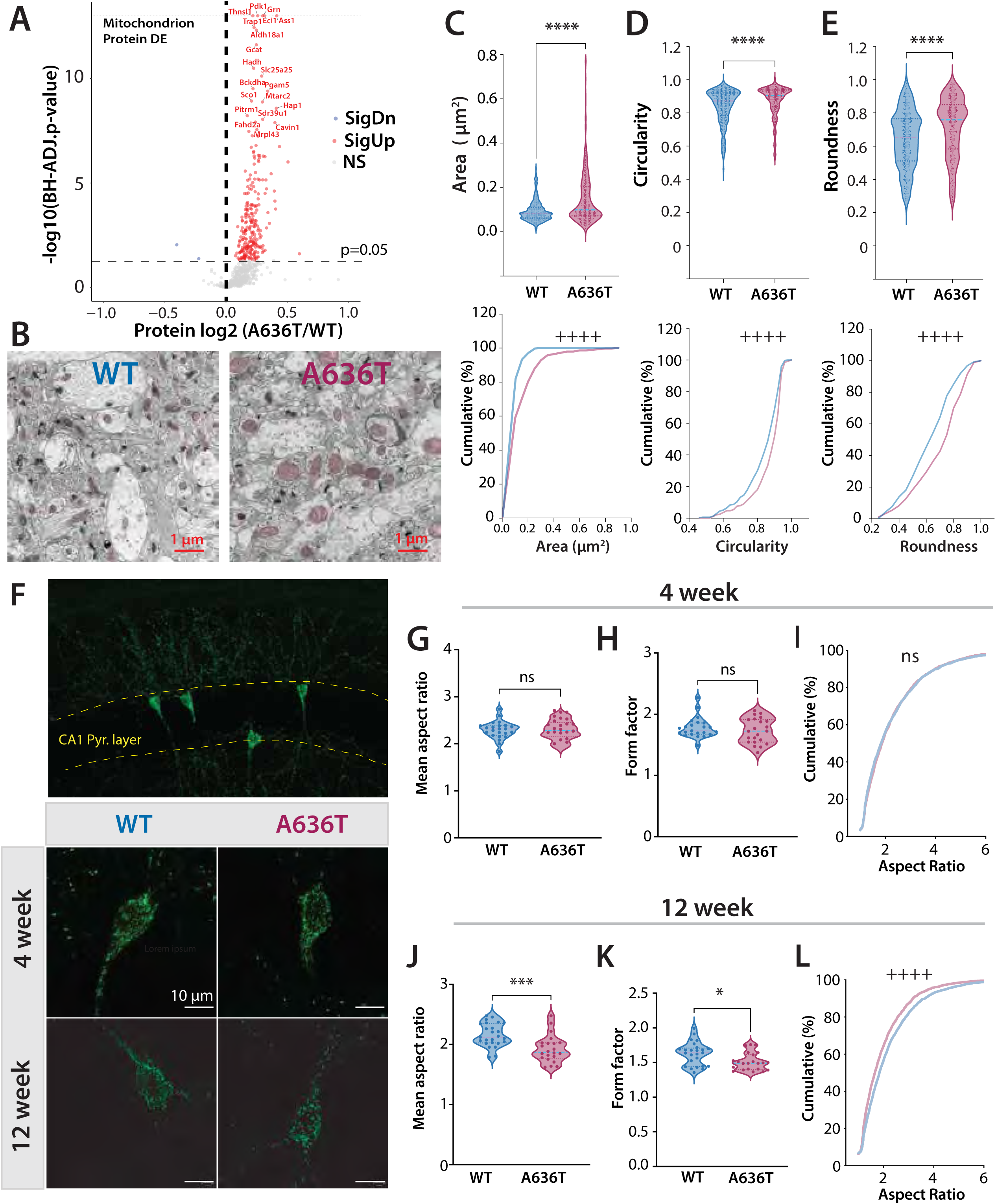
Pathogenic GluA1A636T expression is associated with altered mitochondrial morphology. (**A**) Alterations of mitochondrial proteins in adult WT versus GluA1^A636T^ mouse hippocampus filtered using MitoCarta 3.0. (**B**) Example EM micrograph from stratum radiatum of WT (left) and GluA1^A636T^ mice (right). (**C-E**) (upper panels) Violin plots of analysis of mitochondria; (**C**) area (**D**) circularity (**E**) roundness (lower panels) have cumulative distributions of mitochondria measures. For each animal, 3-5 regions of stratum radiatum about 100–200 μm from the pyramidal cell layer were selected for analysis. Approximately 300 mitochondria per genotype were analyzed (WT: n = 3 mice; GluA1^A636T^: n = 3 mice). ****P < 0.001,Student’s *t*-test. ^++++^P < 0.001, Kolmogorov–Smirnov test. (**F**) Example images of CA1 neurons expressing mito-GFP following RO administration of PHP.eB-hSyn-mito-GFP. Lower panels are example CA1 neurons expressing mito-GFP from WT and GluA1^A636T^ animals at 4 weeks and 12 weeks of age. (**G-I**) Analysis of morphology of mito-GFP expressing mitochondria in 4 week old mice showing mean aspect ratio (**G**) and form factor (**H**). (WT: n = 23 cells from 4 mice; GluA1^A636T^ : n = 22 cells from 3 mice) (**I**) Cumulative plot of aspect ratio of all mitochondira analyzed (approximately 3,000 mitochondria per genotype). (**J-L**) Mitochondrial morphology assessed in 12 week old mice demonstrating significant reduction in mean aspect ratio (**J**) and form factor (**K**) in GluA1^A636T^ mice. (WT: n = 25 cells from 4 mice; GluA1^A636T^: n = 22 cells from 4 mice) *P < 0.05; *** P < 0.001, Student’s *t*-test. (**L**) Cumulative distribution of aspect ratios for all mitochondria analyzed (approximately 3,000 mitochondria per genotype),showing a leftward shift in GluA1^A636T^ mice. ^++++^P < 0.0001 (Kolmogorov–Smirnov test). See Dataset S4 for the detailed MitoCarta 3.0-annotated analysis from the adult TMT-MS dataset.

To assess mitochondrial ultrastructure, we performed transmission electron microscopy (EM) analysis of the stratum radiatum in hippocampal sections from 12-week-old GluA1^A636T^ (n = 3) and littermate control mice (n = 3) (**Figure 3 B**). For each mouse, 3–5 electron micrographs were acquired from the stratum radiatum at a distance of approximately 100-200 μm from the border of the CA1 pyramidal cell layer, yielding a total of approximately 250–300 mitochondria quantified per genotype. Quantitative analysis demonstrated that mitochondria from GluA1^A636T^ mice exhibited significant larger areas (Student’s t test, P < 0.0001; **Figure 3C**), circularity (Student’s t test, P < 0.0001; **Figure 3D**), and roundness (Student’s t test, P < 0.0001; **Figure 3E**), consistent with a shift toward less elongated, more rounded mitochondrial morphology.

To further characterize these morphological changes, we also analyzed the cumulative distributions of individual mitochondrial area, circularity, and roundness (**Figure 3 C-E lower panels)**. The GluA1^A636T^ mice exhibited rightward shifts in the cumulative distributions of mitochondrial area (**Figure 3C**, lower), circularity (**Figure 3D**, lower) , and roundness (**Figure 3E**, lower) (Kolmogorov–Smirnov test, P < 0.0001 for each comparison), indicating a greater proportion of larger and more rounded mitochondria compared with measurements from WT controls.

To determine whether changes in mitochondrial form were also present within identified CA1 pyramidal neurons, as well as to establish when they emerged during development, we next visualized mitochondria using sparse neuronal expression of mitochondria-targeted GFP (mito-GFP; AAV-PHP.eB-hSyn-mito-GFP). Virus was delivered by retro-orbital injection to GluA1^A636T^ mice and littermate controls (Chan et al., 2017) 10-14 days before sacrifice. Mitochondria in soma and proximal dendrites were analyzed in CA1 pyramidal neurons from both juvenile (4-week old) and adult (10-12 week old ) mice (**Figure 3F-L**). The 4-week old group included 4 WT mice and 3 GluA1^A636T^ mice (mixed gender) and the 12-week group include 4 WT mice and 4 GluA1^A636T^ mice (mixed gender). Approximately five CA1 pyramidal neurons were analyzed per mouse. Cell-based morphometric analysis was performed using the Fiji MitochondriaAnalyzer plugin (Chaudhry et al., 2020). In 4 week old mice mitochondrial morphology was indistinguishable between genotypes, with no differences in mitochondrial abundance (Student’s t-test, P = 0.77), mitochondrial area (P = 0.079), aspect ratio (P = 0.861; **Figure 3G**) or form factor (P = 0.363; **Figure 3H**) (WT: 23 cells; GluA1^A636T^: 22 cells ). In adult mice at 12 weeks old similarly mitochondrial abundance (Student’s t-test, P = 0.77) and area (P = 0.91) remained unchanged. However, in contrast to the young mice, adult GluA1^A636T^ mice exhibited a significant reduction in mitochondrial aspect ratio (Student’s t-test, P = 0.00045; **Figure 3J**) and form factor (P = 0.011; **Figure 3K**) (WT: 25 cells; GluA1^A636T^: 22 cells) indicating a shift toward a rounder and less elongated mitochondria morphology.

To provide a more comprehensive assessment of variability, we analyzed the cumulative distribution of mitochondrial aspect ratios by pooling mitochondria from all cells of the same genotype. At 4 weeks of age the cumulative distributions of aspect ratio did not differ significantly between WT and GluA1^A636T^ mice (Kolmogorov–Smirnov test, P = 0.0535; **Figure 3I**)(approximately 3,000 mitochondria per genotype). In contrast, at 12 weeks of age, the cumulative aspect ratio distributions differed significantly between genotypes, with the GluA1^A636T^ mice exhibiting a leftward shift indicating a greater proportion of less elongated mitochondria (Kolmogorov–Smirnov test, P < 0.0001; **Figure 3L**) (approximately 3,000 mitochondria per genotype). Together, these complementary analyses demonstrate that pathogenic GluA1 A636T expression is associated with an age-dependent remodeling of mitochondrial morphology that emerges during adulthood without a detectable change in mitochondrial abundance.

### Early oxidative stress precedes mitochondrial remodeling in GluA1^A636T^ mice

Because proteomic analysis identified alterations in proteins associated with oxidative stress, we next examined markers of oxidative damage in the hippocampus across development. Immunohistochemical analysis of the oxidative and nitrosative stress marker 3-nitrotyrosine (3NT) and the reactive oxygen species indicator dihydroethidium (DHE) revealed increased signal in GluA1^A636T^ mice at both juvenile (4-week) and adult (12-week) ages compared with littermate controls (**Figure 4 A-D**) Two-way ANOVA revealed a significant main effect of genotype for both 3NT immunoreactivity (genotype: F_1,13_ = 166.83, P < 0.0001)(**Figure 4B**) and DHE staining (genotype: F_1,18_ = 151.23, P < 0.0001)(**Figure 4D**) (3NT staining: 4-week group, 5 WT and 4 GluA1^A636T^ mice; 12-week group, 4 WT and 4 GluA1^A636T^ mice) (DHE staining: 4-week group, 5 WT and 5 GluA1^A636T^ mice; 12-week group, 5 WT and 6 GluA1^A636T^mice). There was no significant genotype × age interaction for either marker (3NT: F_1,13_ = 1.29, P = 0.277; DHE: F_1,18_ = 0.22, P = 0.644), indicating that oxidative stress is already elevated during juvenile development and persists into adulthood. Notably, this temporal profile differs from the mitochondrial morphological phenotype, which was not detected at 4 weeks but became evident by 12 weeks. (**Figure 3**). Together, these findings demonstrate that pathogenic GluA1A636T expression is associated with oxidative stress that is already evident during juvenile development and persists into adulthood, whereas detectable remodeling of mitochondrial morphology emerges later.

**Figure 4.**
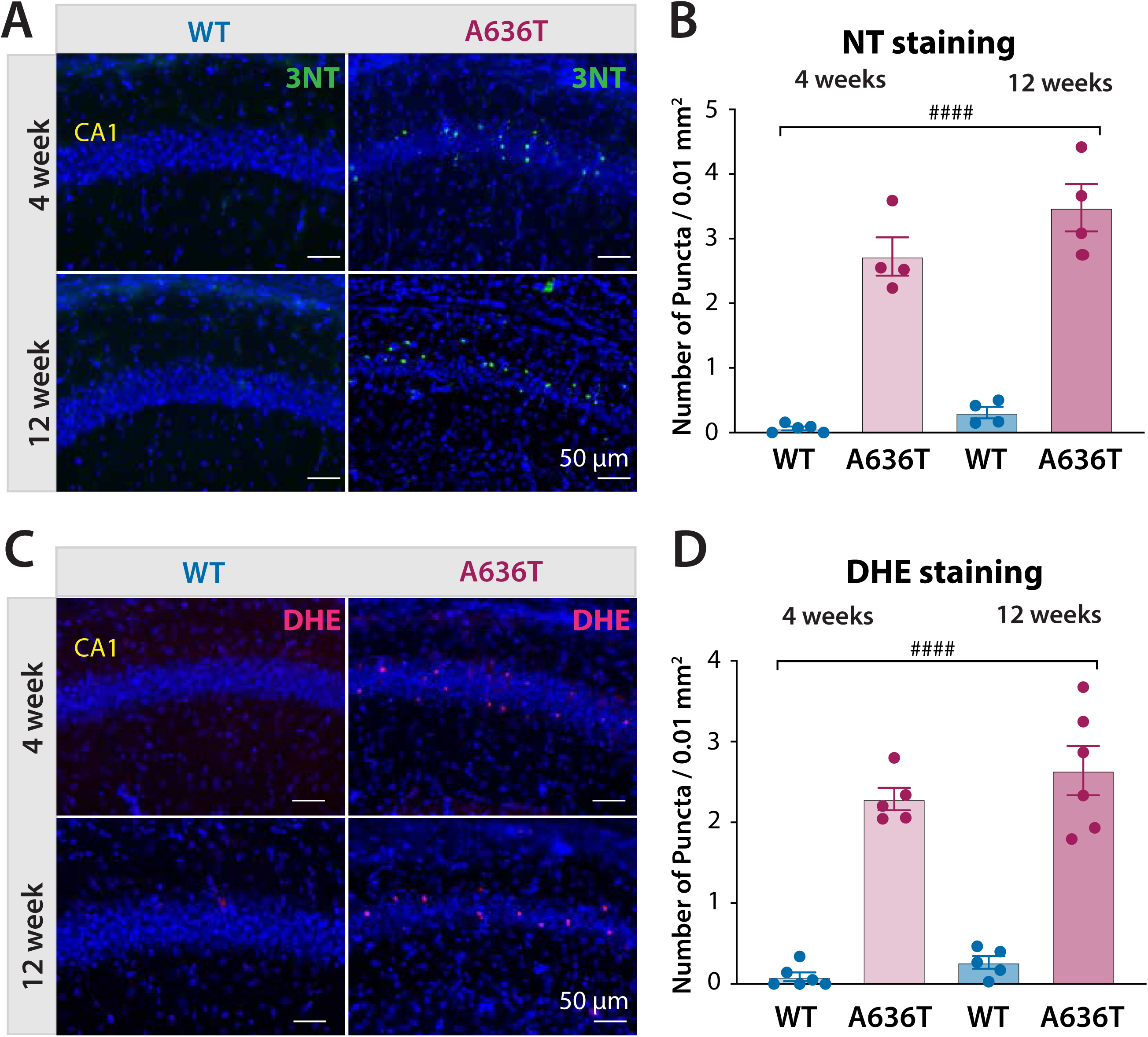
Increased oxidative stress markers in the hippocampus of GluA1^A636T^ mice. (**A**) Example images of staining in WT and GluA1^A636T^ mice of 3-nitrotyrosine (NT) in 4 week old and 12 week old mice and (**B**) Analysis of NT staining in mice of each genotype at juvenile and adult ages (**C**) Example images of immunohistochemical staining in WT and GluA1^A636T^ of dihydroethidium (DHE) (**D**) Analysis of DHE staining in WT and GluA1^A636T^ mice at 4 week and 12 weeks of age. ^####^ P < 0.0001, Two-way ANOVA. Data are represented as mean ± SEM. Error bars represent SEM.

### Basal excitatory synaptic function is disrupted in GluA1^A636T^ hippocampus

Proteomic analysis identified alterations in numerous synaptic proteins at both juvenile and adult ages, suggesting that pathogenic GluA1A636T expression disrupts the molecular organization of excitatory synapses. We therefore examined hippocampal synaptic transmission and plasticity in acute brain slices from juvenile (3–4 weeks) and adult (10–12 weeks) mice to determine whether functional deficits exist in CA1 synapses during postnatal development.

To assess basal excitatory synaptic transmission, mEPSCs were recorded from CA1 pyramidal neurons (**Figure 5A -D**). We found that the decay time (τ) of AMPAR mediated mEPSCs was significantly increased in juvenile GluA1^A636T^ mice (p < 0.0001, Mann-Whitney, 21 cells from 3 WT mice, 22 cells from 4 GluA1^A636T^ mice) (**Figure 5B, left**). This finding is consistent with previous work reporting that the GluA1A636T mutation slows the deactivation kinetics of AMPARs (Kohda et al., 2000; Klein and Howe, 2004), and suggests that A636T mutant GluA1 subunits are incorporated into functional AMPARs *in vivo*. We also found a significant decrease in the frequency of mEPSCs in juvenile GluA1^A636T^ mice (p=0.0038, Mann-Whitney, 21 cells from 3 WT mice, 22 cells from 4 GluA1^A636T^ mice) (**Figure 5B, center)** while mEPSC amplitudes were larger in the 3-4 week old mutant mice (p = 0.011, Mann-Whitney, 21 cells from 3 WT mice, 22 cells from 4 GluA1^A636T^ mice) (**Figure 5B, right**).

**Figure 5.**
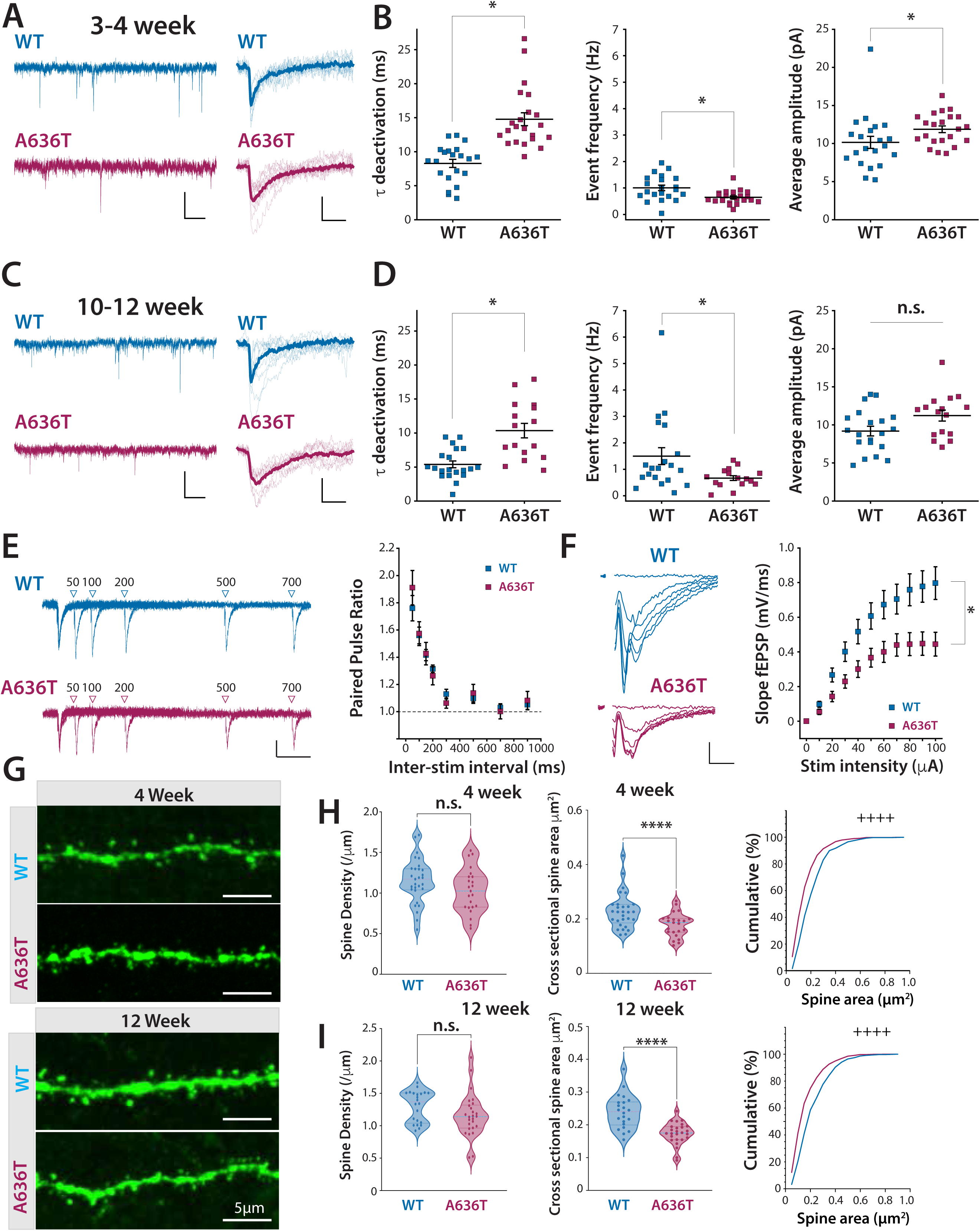
Basal excitatory synaptic transmission and spine density in CA1 of GluA1^A636T^ hippocampus. (**A**) Representative mEPSC traces from juvenile mice (3–4 weeks). Calibration: 10 pA, 1 s, and individual events with the average of 10 events in bold. Calibration: 5 pA, 10 ms (**B**) Summary quantification of mEPSC decay time constant (τ; left), event frequency (Hz; middle), and mean amplitude (pA; right) in juvenile mice. (**C**) Representative mEPSC traces from adult (10-12 week) mice Calibration: 10 pA, 1 s and individual events shown with the average of 10 representative mEPSCs in bold. Calibration: 5 pA, 10 ms (**D**) Summary quantification of mEPSC decay time constant (τ; left), event frequency (Hz; middle), and mean amplitude (pA; right) in adult mice. *p < 0.05. (**E**) Representative CA1 fEPSP traces evoked by paired stimuli delivered at varying interstimulus intervals (ISIs) indicated by arrowheads indicate Calibration: 0.2 mV, 100 ms. Right panel shows summary of paired-pulse ratio measured in CA1 (WT: n = 24 cells from 8 mice; GluA1^A636T^: n = 15 cells from 4 mice). (**F**) Representative fEPSP traces evoked by incrementally increasing stimulation intensity. Calibration: 0.2 mV, 5 ms. Right panel shows summary of input–output relationship of SC-CA1 EPSP (WT: n = 24 cells from 8 mice; GluA1^A636T^: n = 22 cells from 5 mice). *P < 0.05. (**G**) Example images of dendritic segments from WT and GluA1^A636T^ mice at 4 weeks and 12 weeks of age. (**H**) Quantification of spine density (**left panel**) and mean cross sectional spine area (**middle panel**) (average per dendritic segment) in 4-week-old mice. The **right panel** shows cumulative distribution of spine areas of all measured spines (WT: n = 29 segment from 4 mice; GluA1^A636**T**^: 23 from 4 mice. Cumulative spine area distributions were generated from approximately 500 spines per genotype. (**I**) Quantification of spine density (**left panel**) and mean cross-sectional spine area (**middle panel**) (average per dendritic segment) in 12-week-old mice. The **right panel** shows the cumulative distribution of spine areas for all measured spines (WT: n = 21 from 4 mice; GluA1^A636**T**^: n = 25 from 4 mice; Cumulative spine area distributions were generated from approximately 600 spines per genotype. *P < 0.05; ****P < 0.0001(Student’s *t*-test); ^++++^P < 0.0001 (Kolmogorov–Smirnov test).

In adult mice (10-12 wks), we observed similar synaptic alterations where the decay τ was slower (P < 0.0001, Mann-Whitney, 20 cells from 5 WT mice, 16 cells from 4 GluA1^A636T^ mice) (**Figure 5 C, D left**), the frequency was decreased (p = 0.037, Mann-Whitney, 20 cells from 5 WT mice, 16 cells from 4 GluA1^A636T^ mice) (**Figure 5D, middle**), while comparisons of the amplitudes were not significant (P = 0.059, Mann-Whitney, 20 cells from 5 WT mice, 16 cells from 4 GluA1^A636T^ mice) (**Figure 5D, right**) in GluA1^A636T^ mice.

Reduced mEPSC frequency can reflect either a decrease in the number of functional synapses or a reduction in presynaptic release probability (RP). To distinguish between these possibilities, we measured paired pulse ratio (PPR) of field EPSPs (fEPSPs) at Schaffer collateral – CA1 synapses in adult mice. PPR was unchanged in GluA1^A636T^ mice compared to WT controls (**Figure 5 D**), arguing against a major alteration in presynaptic release probability.

We next examined the input–output (I-O) relationship of CA1 fEPSPs which reflects how the postsynaptic response increases with progressively stronger presynaptic stimulation. GluA1^A636T^ mice exhibited a significant downward shift in the I-O curves in relation to recordings from WT littermates (**Figure 5 F)** indicating reduced postsynaptic responsiveness and diminished synaptic recruitment across stimulation intensities. Taken together with the reduction in mEPSC frequency and unchanged PPR, these findings are most consistent with a decrease in the number of functionally active synapses in GluA1^A636T^ mice rather than impaired release probability. This interpretation is further supported by the observed downregulation of synaptic protein levels in the hippocampus (**Figure 2**), collectively indicating weakened excitatory connectivity in GluA1^A636T^ mice.

Because synaptic deficits could reflect either a loss of excitatory contacts or altered structural maturation of existing synapses, we next examined whether the functional abnormalities in excitatory transmission were accompanied by alterations in dendritic spine morphology. Our analysis focused on second order apical dendrites of CA1 pyramidal neurons. At the 4 week old juvenile time point, spine density did not differ significantly between WT and GluA1^A636T^ mice (WT: 1.151 ± 0.051 spines/µm, 29 dendritic segments from 4 mice; GluA1^A636T^: 1.029 ± 0.057 spines/µm, 23 dendritic segments from 4 mice; unpaired two-tailed Student’s *t*-test, P = 0.1146) (**Figure 5H, left**). In contrast, mean spine cross-sectional area was significantly reduced in GluA1^A636T^ mice (WT: 0.229 ± 0.012 µm², 29 dendritic segments from 4 mice; GluA1^A636T^: 0.175 ± 0.0048 µm², 23 dendritic segments from 4 mice; P < 0.0001) (**Figure 5H, middle**). The distribution of individual spine areas was also shifted toward smaller values in mutant neurons ( > 500 spines from each genotype, Kolmogorov–Smirnov test, P < 0.0001), indicating a reduction in spine size (**Figure 5H, right**).

We next asked whether these structural alterations persisted into adulthood (**Figure 5G, I**). Similar to juvenile mice, spine density of CA1 secondary apical dendrites did not differ between genotypes at 12 weeks of age (**Figure 5I, left**) (WT: 1.303 ± 0.051 spines/µm, 22 dendritic segments from 4 mice; GluA1^A636T^: 1.157 ± 0.066 spines/µm, 26 dendritic segments from 4 mice; Student’s *t*-test, *P* = 0.097), whereas spine cross-sectional area remained significantly reduced in GluA1^A636T^ mice compared to WT (**Figure 5I, middle**) (WT: 0.240 ± 0.0051 µm², 22 dendritic segments; GluA1^A636T^: 0.174 ± 0.0335 µm², 26 dendritic segments; P < 0.0001). Again, the cumulative distribution of individual spine areas was shifted toward smaller values in mutant neurons (Kolmogorov–Smirnov test, P < 0.0001)(approximately 600 spines per genotype; **Figure 5I, right**), consistent with reduced spine size.

### NMDAR-only silent synapses remain prevalent in adulthood in GluA1^A636T^ mice

Analyses of basal synaptic transmission together with the reduction in dendritic spine size suggested that a subset of excitatory contacts may remain structurally present but functionally immature in the adult GluA1^A636T^ hippocampus. One potential explanation is that these synapses fail to acquire stable AMPAR-mediated transmission and therefore persist as NMDAR-only silent synapses. To directly test this possibility we next examined the prevalence of NMDAR-only synapses in adult GluA1^A636T^ mice.

Assessment of the presence of silent synapses was made in 10-12-week-old mice when under normal conditions, they are undetectable in the hippocampus based on functional measures (Hanse et al., 2013). As the primary measure of silent synapses we employed a minimal stimulation protocol designed to recruit single or a small number of presynaptic axons and measured the failure rate of evoked unitary synaptic responses (Liao et al., 1995; Harlow et al., 2010) (**Figure 6 A-E**). EPSCs were stimulated while holding the cell at -70 mV to record the AMPAR-mediated component and +40 mV to record the mixed AMPA and NMDAR-mediated component of the EPSCs (**Figure 6 A & B**). The failure rate at each holding potential was calculated in each recorded neuron (**Figure 6 C & D**). From this we calculated the failure ratio as the failure rate at +40mV divided by the failure rate at -70mV (**Figure 6 E**). In recordings from WT controls the failure ratio was close to 1 suggesting that there were few detectable silent synapses in WT adult mice as previously reported (**Figure 6 E**) (Isaac et al., 1995; Liao et al., 1995; Durand et al., 1996). In the GluA1^A636T^ mice we detected a higher failure rate at a holding potential of -70 mV compared to +40 mV, with a +40/-70 mV failure ratio lower than 1 on average demonstrating that there is a likely higher retention of silent synapses (**Figure 6 E**). We also performed a quantitative assessment of this data using previous methods (Liao et al., 1995) and estimated that in control mice there were 12.5% abundance of silent synapses, while in GluA1^A636T^ mice we calculated an abundance estimate of 24.6% silent synapses.

**Figure 6.**
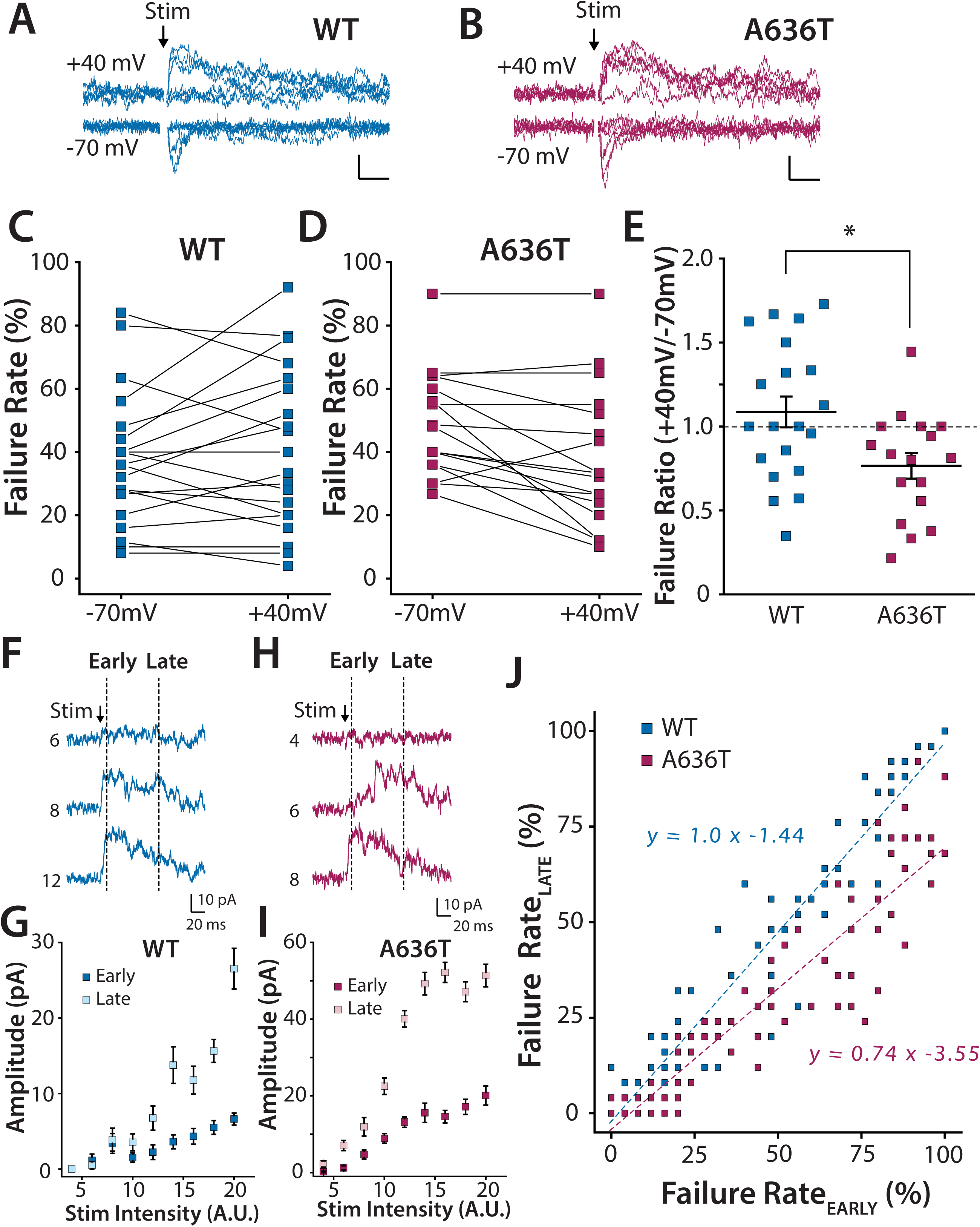
Persistent silent synapses in adult GluA1^A636T^ hippocampus. (**A**) Representative minimally evoked EPSCs and failures recorded from WT neurons at +40 mV and −70 mV (six traces per holding potential). (**B**) Representative minimally evoked EPSCs and failures recorded from GluA1^A636T^ mice. Calibration; 10pA, 20ms (**C**) Failure rate of EPSCs at −70 mV and +40 mV in WT mice (**D**) EPSC failure rate at −70 mV and +40 mV in GluA1^A636T^ mice. (WT: n = 20 cells from 6 mice; GluA1^A636T^: n = 17 cells from 5 mice). (**E**) Failure ratio (+40 mV / −70 mV) for all recordings (Wilcoxon test, *p < 0.05). (**F**) Representative EPSCs recorded at +40 mV using a modified minimal stimulation protocol in WT mice. Stimulation intensities (A.U.) are indicated. Early (0.5–1 ms post-stimulus) and late (50–50.5 ms post-stimulus) EPSC components are indicated by dashed windows. Calibration: 10 pA, 20 ms (**G**) Input–output (I-O) relationship of stimulation intensity and mean amplitude of early and late EPSC components (EPSC_EARLY_ and EPSC_LATE_) for WT recording. Mean amplitudes from 25 sweeps (including failures) are plotted at each stimulation intensity. (**H**) Representative EPSCs recorded with same protocol in GluA1^A636T^ mice. Calibration: 10 pA, 20 ms (**I**) I-O relationship of EPSC_EARLY_ and EPSC_LATE_ for GluA1^A636T^ recordings. (**J**) Summary plot of failure rates of EPSC_LATE_ versus EPSC_EARLY_ across all stimulation intensities and recorded cells in WT and GluA1^A636T^ mice. Significant deviation from the unity line is observed in GluA1^A636T^ mice but not in WT controls. *P < 0.05.

As a secondary empirical measure to compare the abundance of silent synapses in GluA1^A636T^ mice, we performed a complementary analysis using an incremental stimulation method to record the early and late components of the EPSC (EPSC_E_ and EPSC_L_; see Methods) while holding the neuron at a membrane potential of +40 mV to enable detection of both AMPAR-and NMDAR-mediated EPSCs (Isaac et al., 1995). For each stimulation intensity, we quantified the presence or absence of early (predominantly AMPA mediated) and late EPSC (predominantly NMDA mediated) components (**Figure 6F-I**). In WT recordings, both EPSC_E_ and EPSC_L_ emerged concomitantly as stimulus intensity increased (**Figure 6 F & G**). In contrast, in GluA1^A636T^ mice, EPSC_L_ responses were frequently observed in the absence of detectable EPSC_E_ (**Figure 6 H & J**). This was also evident in the trend lines when plotting failure rates of EPSC_LATE_ against EPSC_EARLY_ for all recordings where the trend line was close to unity in WT recordings but below unity in GluA1^A636T^ recordings (**Figure 6 J**) consistent with an increased prevalence of synapses that contain NMDARs but lack functional AMPAR-mediated transmission.

Taken together, these findings indicate that there is an elevation of NMDAR only synapses in adult GluA1^A636T^ mice and suggests that their normal developmental elimination (Durand et al., 1996) is incomplete in GluA1^A636T^ mice. Persistent retention of silent synapses into adulthood is therefore likely to contribute, in part, to the reduced functional synaptic connectivity observed in the mutant hippocampus.

### Activity-dependent un-silencing and enhanced potentiation in GluA1^A636T^ mice

Silent synapses are not inert but rather represent a population of developmentally immature connections that are particularly susceptible to activity-dependent un-silencing through AMPAR recruitment. The increased abundance of NMDAR-only synapses observed in GluA1^A636T^ mice therefore predicts an altered capacity for synaptic potentiation when appropriate patterns of activity are imposed. To test whether aberrant retention of silent synapses translates into altered synaptic plasticity at the circuit level, we next examined if LTP inducing stimuli differentially affect in WT and GluA1^A636T^ mice.

We assessed activity-dependent unsilencing of silent synapses at SC–CA1 synapses using a minimal stimulation protocol. After establishing a stable baseline for 5 minutes, brief train stimulation (50 pulses at 10 ms intervals) was delivered to induce long-term potentiation (LTP). In GluA1^A636T^ mice, train stimulation produced a significant reduction in EPSC failure rate, consistent with recruitment of previously silent synapses (**Figure 7A–D**). In contrast, the same stimulation paradigm produced little or no change in failure rate in WT mice (**Figure 7D**).

**Figure 7.**
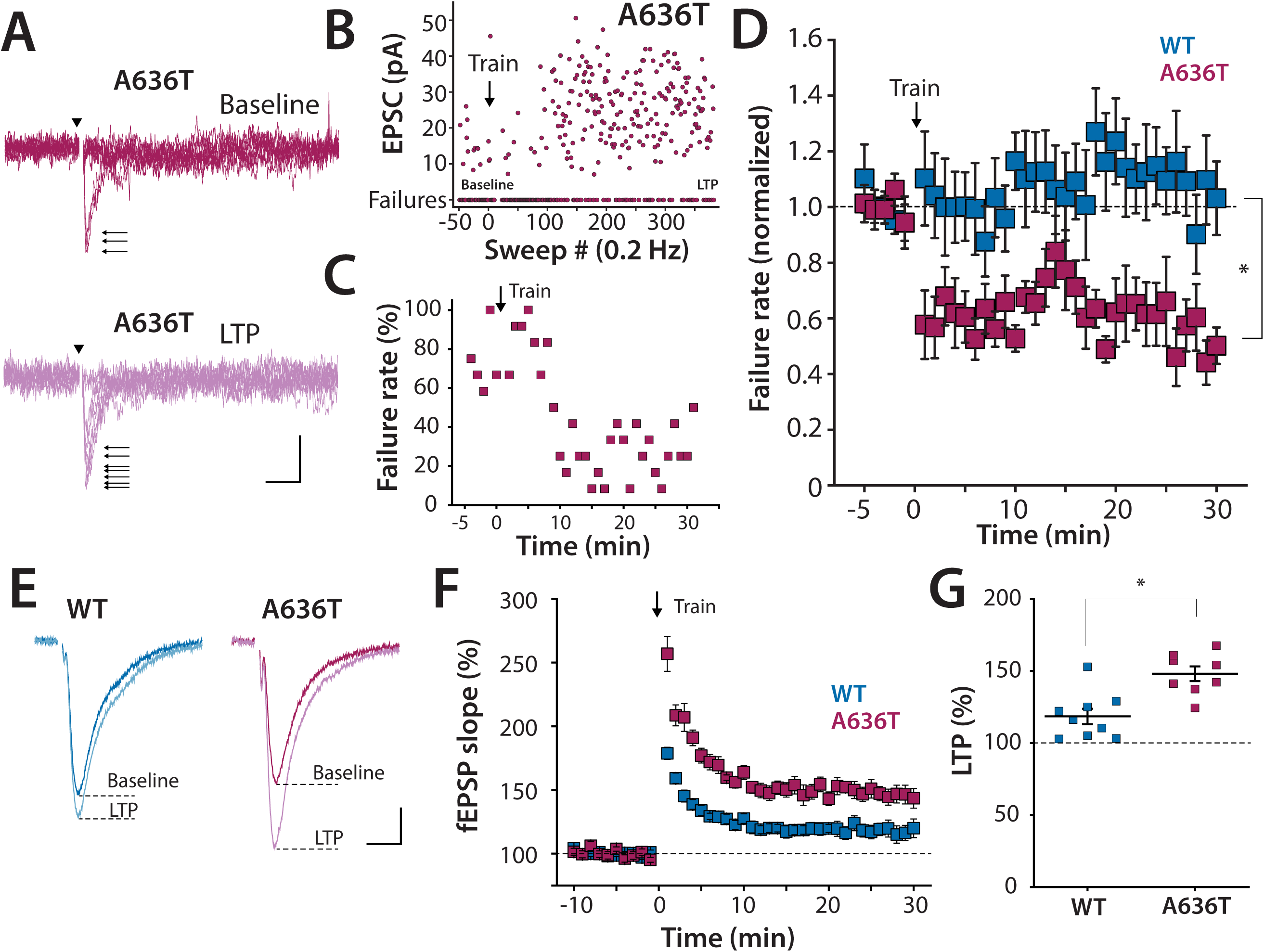
Activity-dependent unsilencing is associated with enhanced synaptic plasticity in GluA1^A636T^ hippocampus. (**A**) Representative minimally evoked EPSCs (arrows) and failures recorded at baseline and following high frequency stimulation in GluA1^A636T^ mice (10 traces per condition). Arrowheads indicate delivery of minimal stimuli. Scale bars: 20 ms and 10 pA. (**B**) Example time course of synaptic responses and (**C**) failure rate during the unsilencing paradigm in a single example recording. (**D**) Summary of normalized failure rates for all recordings before and after unsilencing in WT and GluA1^A636T^ mice. (**E**) Example fEPSP traces in WT and GluA1^A636T^ mice before and after LTP induction. Calibration: 10 ms and 0.1 mV. (**F**) Time course of LTP induced by brief high frequency stimulation. (**G**) Summary of LTP magnitude in all recordings. *P < 0.05.

Together, these results indicate that silent synapses are more prevalent in adult GluA1^A636T^ mice and are readily unsilenced by LTP-inducing activity, whereas silent synapses are largely absent or rare in WT hippocampus at this developmental stage.

We next asked whether enhanced synaptic unsilencing is associated with an increased capacity for activity-dependent synaptic potentiation in GluA1^A636T^ mice. To test this, we applied the same train stimulation paradigm used for unsilencing experiments to SC–CA1 synapses in an extracellular field recording configuration. In WT slices, this stimulation reliably induced modest long-term potentiation (∼120% of baseline). In contrast, GluA1^A636T^ slices exhibited significantly greater potentiation (∼150% of baseline) (**Figure 7 E-G**). These findings indicate that persistent silent synapses in adult GluA1^A636T^ mice lower the effective threshold for activity-dependent synaptic strengthening, resulting in exaggerated potentiation when unsilencing mechanisms are engaged.

Collectively, these finding demonstrate persistent reductions in synaptic transmission, smaller dendritic spines, retention of NMDAR-only silent synapses, and enhanced activity-dependent potentiation which together indicate that many excitatory contacts fail to complete the normal transition to mature AMPAR-rich synapses and thus identify impaired developmental maturation of excitatory synapses as a central consequence of pathogenic GluA1 dysfunction.

## Discussion

In this study, we demonstrate that a recurrent pathogenic human mutation at the conserved Lurcher site of the AMPA receptor GluA1 subunit (Geisheker et al., 2017) disrupts the coordinated maturation of excitatory synapses and causes cellular stress responses in the mouse hippocampus. By integrating developmental proteomics and transcriptomics with synaptic physiology, and behavior, we show that GluA1^A636T^ expression leads to a reduction in functional excitatory connectivity, persistent retention of silent synapses in adulthood, and altered activity-dependent unsilencing, accompanied by progressive oxidative stress and delayed mitochondrial remodeling. Together these findings reveal that disease-associated AMPAR mutations can impair brain function not through uniform changes in synaptic strength, but by perturbing the temporal alignment of synapse maturation and metabolic support mechanisms that normally stabilize developing circuits (Katz and Shatz, 1996)

The A636T substitution lies within the highly conserved SYTANLAAF motif of the M3 transmembrane helix, commonly referred to as the Lurcher site, which plays a central role in channel gating in ionotropic glutamate receptors (Klein and Howe, 2004; Sobolevsky et al., 2009). Missense mutations at this site recur across multiple glutamate receptor subfamilies, including AMPA, NMDA, and kainate receptors, and are disproportionately associated with neurodevelopmental disorders (Davies et al., 2017; Guzman et al., 2017). While such variants are often classified as gain-of-function based on altered channel kinetics in heterologous systems, our findings underscore that such biophysical classifications alone are insufficient to predict in vivo consequences, which instead reflect complex developmental interactions between synaptic signaling, circuit refinement, and cellular homeostasis (Myers et al., 2023; Nomura et al., 2023).

A central insight gained from our study is that the primary consequence of GluA1A636T expression is the disruption of synapse maturation rather than a uniform alteration in synaptic efficacy. During normal development, activity-dependent recruitment of AMPARs drives the conversion of NMDAR-only synapses into fully functional excitatory contacts, stabilizing emerging circuits and constraining plasticity in adulthood (Durand et al., 1996; Isaac et al., 1997; Pickard et al., 2000). In adult GluA1^A636T^ mice, we instead observe a persistent population of functionally silent synapses, reduced basal excitatory connectivity, and exaggerated activity-dependent un-silencing. This constellation of phenotypes is inconsistent with a simple gain-or loss-of-function model and instead indicates that pathogenic AMPAR subunit mutation can arrest or delay the normal developmental trajectory of excitatory synapse refinement. Importantly, the present study does not establish how the A636T mutation alters the biophysical properties, subunit composition, trafficking, or synaptic stabilization of native heteromeric AMPARs. Thus, while our data establish the consequences of A636T expression for synaptic development and function in vivo, the molecular mechanism linking the mutant subunit to impaired synapse maturation remains to be determined.

The enhanced potentiation observed in adult GluA1^A636T^ mice does not reflect strengthened synaptic transmission per se, but rather the recruitment of synapses that failed to mature appropriately during development. Such retained developmental plasticity rules in adult circuits have been observed when early activity-dependent processes are perturbed and are often accompanied by impaired information processing and behavioral deficits despite preserved or even exaggerated synaptic plasticity (Katz and Shatz, 1996; Turrigiano and Nelson, 2004; Hensch, 2005). Viewed in this context, the increased capacity for synaptic un-silencing in GluA1^A636T^ mice represents a maladaptive persistence of immature circuit states, linking disrupted synapse maturation to long-lasting alterations in hippocampal function.

Beyond synaptic maturation, our data also reveals a striking temporal dissociation between early synaptic alterations and later mitochondrial remodeling, pointing to a hierarchical relationship between excitatory signaling and cellular metabolic homeostasis. Proteomic profiling across development showed that reductions in synaptic protein abundance emerge prior to detectable changes in mitochondrial pathways, whereas in adult GluA1^A636T^ mice mitochondrial proteins are selectively upregulated in a manner largely uncoupled from transcriptional regulation. This proteome–transcriptome divergence, together with the absence of mitochondrial pathway enrichment at earlier developmental stages, argues against a primary mitochondrial defect and instead suggests that early synaptic alterations and later mitochondrial remodeling represent distinct but potentially related components of the evolving phenotype.

Our integrated proteomic and transcriptomic analyses also reveal that signatures of oxidative stress, inflammatory signaling, and apoptosis-related pathways are already present at the juvenile stage, prior to the emergence of robust mitochondrial protein changes in GluA1^A636T^ mice. These early signals are further supported by independent measurements of markers of oxidative stress in the hippocampus in juveniles, indicating that cellular stress responses are engaged early in development alongside synaptic alterations. In adult animals, these pathways become more pronounced and are accompanied by a marked expansion of glial-associated and mitochondrial protein signatures in the adult hippocampus. This temporal pattern suggests that early synaptic disruption is associated with an initial activation of stress and inflammatory responses that intensify over development and coincide with broader metabolic and cellular remodeling in adulthood, with mitochondrial alterations emerging as part of this progression.

Consistent with these molecular changes, mitochondria in GluA1^A636T^ hippocampus exhibited altered morphology in adult mice but not at juvenile ages, and increased markers of oxidative stress at both ages; phenotypes that have been linked to sustained changes in neuronal activity and cellular demand rather than acute excitotoxic injury (Li et al., 2004; Harris et al., 2012; Datta and Jaiswal, 2021; Virga et al., 2024). Mitochondria play a central role in supporting cellular energy demands, calcium buffering, and metabolic homeostasis and their structure and function are sensitive to changes in neuronal activity (Virga et al., 2024). Together these observations indicate that mitochondrial alterations are a prominent feature of the adult phenotype and are associated with the broader molecular and cellular changes across development.

The synaptic abnormalities identified here would be expected to alter the fidelity of information processing within hippocampal circuits rather than simply suppress overall neuronal activity. Although CA1 pyramidal neurons in GluA1^A636T^ mice exhibited activity levels comparable to WT animals across locomotor states, this does not imply normal circuit function. Stable acquisition of AMPAR-mediated transmission is critical for the maturation and refinement of hippocampal ensembles that encode spatial and contextual information. Interestingly CA1 neurons in GluA1^-/-^ mice maintain normal firing rates despite impaired spatial coding, suggesting that overall neuronal activity can be preserved even when hippocampal information processing is disrupted (Resnik et al., 2012). Persistent silent synapses, smaller dendritic spines, and reduced functional excitatory transmission would therefore be predicted to compromise the precision or stability of these representations even if overall neuronal activity remains intact.

Consistent with this interpretation, GluA1^A636T^ mice exhibited impairments in both spatial learning and working memory, demonstrating that disruption of synaptic maturation is accompanied by deficits in hippocampal-dependent cognition. While our calcium imaging did not reveal large changes in overall CA1 activity during locomotion, the analysis was not designed to assess how spatial or mnemonic information is represented within neuronal ensembles. Future experiments examining place cell stability, spatial coding, or ensemble dynamics will be required to determine how impaired synapse maturation alters hippocampal information processing in vivo. Together, these findings suggest that impaired developmental maturation of excitatory synapses is sufficient to disrupt hippocampal-dependent behavior despite relatively preserved overall CA1 activity, highlighting the importance of synaptic composition and maturation for the integrity of hippocampal information processing.

The broader implication of this work is that pathogenic variation within conserved elements of ionotropic glutamate receptors may preferentially disrupt developmental coordination. The recurrence of disease-associated mutations within the Lurcher motif across members of the iGluR subfamilies suggests that this region constitutes a shared molecular constraint through which diverse receptors shape circuit assembly and refinement. Our findings add to growing evidence that the functional consequences of such variants depend not only on altered channel properties, but on how those alterations interact with activity-dependent developmental programs, homeostatic regulation, and circuit-specific demands over time. While this work was under preparation, an preprint reported the independent generation of a GluA1^A636T^ knock-in mouse model and identified concordant behavioral phenotypes and synaptic abnormalities, together with evidence for excitotoxic neurodegeneration and therapeutic rescue using allele-specific antisense oligonucleotides (Chen et al., 2025). Our study is distinct in scope, emphasizing developmental synapse maturation, proteome–transcriptome remodeling, mitochondrial alterations, and altered physiology rather than degeneration or rescue. Viewed together, these findings highlight that pathogenic AMPAR mutations can engage multiple pathological processes whose relative contributions may depend on developmental timing and cellular vulnerability.

These findings have important implications for GRI disorders. They suggest that phenotypic heterogeneity among patients may arise not only from differences in receptor biophysical properties, but also from when and how pathogenic variants disrupt the developmental maturation of excitatory synapses. Our results further imply that therapeutic strategies aimed solely at normalizing receptor function in the mature brain may be insufficient to reverse abnormalities that become established during early development. Instead, interventions delivered during critical developmental windows that preserve normal synapse maturation or that limit the emergence of secondary pathological processes such as oxidative stress and mitochondrial dysfunction, may offer greater potential to alter disease progression. More broadly, our study highlights the importance of considering developmental timing and downstream pathological adaptations when investigating disease mechanisms and designing therapeutic strategies for neurodevelopmental disorders caused by pathogenic glutamate receptor variants.

## Acknowledgements

This work was supported by R01 MH130428 to AC and JNS, R01 MH099114 to AC, R01 NS115471 to AC, R01 EY030169 to YZ, R01 EY032506 to AC and YZ, R01 AG078796 and S10 OD032464 to JNS. We thank the Northwestern University NUSeq core facility and the Center for Advanced Microscopy and Nikon imaging at Northwestern for their technical support.

## Author Contributions

JX, YZW, TN, YZ, JNS, AC designed the experiments. JX, YZW, TN, HV, YZ, CCMC, JJM, JNA, YZ performed experiments and analyzed data. JX, YZW, TN, JNS, AC wrote the initial draft of the manuscript.

## Notes

### Competing Interest Statement

The authors have declared no competing interest.

### Summary of Updates

The manuscript was revised after a first round of reviews at Journal of Neuroscience. New data was added and figures were revised to satisfy the concerns of the reviewers. This new version comports with the version that was sent to the journal after review

## Reference

Anitha A, Thanseem I, Iype M, Thomas SV (2023) Mitochondrial dysfunction in cognitive neurodevelopmental disorders: Cause or effect? Mitochondrion 69:18–32.

Block ML, Zecca L, Hong JS (2007) Microglia-mediated neurotoxicity: uncovering the molecular mechanisms. Nat Rev Neurosci 8:57–69.

Chan KY, Jang MJ, Yoo BB, Greenbaum A, Ravi N, Wu WL, Sanchez-Guardado L, Lois C, Mazmanian SK, Deverman BE, Gradinaru V (2017) Engineered AAVs for efficient noninvasive gene delivery to the central and peripheral nervous systems. Nat Neurosci 20:1172–1179.

Chaudhry A, Shi R, Luciani DS (2020) A pipeline for multidimensional confocal analysis of mitochondrial morphology, function, and dynamics in pancreatic beta-cells. Am J Physiol Endocrinol Metab 318:E87–E101.

Chen CM, Huang YM, Chung CC, Johnson RC, Tsai CY, Chen YH, Tan HL, Hsiao FY, Huganir RL, Chiu SL (2025) Rescuing Neurodevelopmental Deficits in AMPA Receptor Gain-of-Function Mutant. bioRxiv.

Chen Y, Qin C, Huang J, Tang X, Liu C, Huang K, Xu J, Guo G, Tong A, Zhou L (2020) The role of astrocytes in oxidative stress of central nervous system: A mixed blessing. Cell Prolif 53:e12781.

Datta S, Jaiswal M (2021) Mitochondrial calcium at the synapse. Mitochondrion 59:135–153.

Davies B et al. (2017) A point mutation in the ion conduction pore of AMPA receptor GRIA3 causes dramatically perturbed sleep patterns as well as intellectual disability. Hum Mol Genet 26:3869–3882.

Dobin A, Davis CA, Schlesinger F, Drenkow J, Zaleski C, Jha S, Batut P, Chaisson M, Gingeras TR (2013) STAR: ultrafast universal RNA-seq aligner. Bioinformatics 29:15–21.

Dombeck DA, Harvey CD, Tian L, Looger LL, Tank DW (2010) Functional imaging of hippocampal place cells at cellular resolution during virtual navigation. Nat Neurosci 13:1433–1440.

Durand GM, Kovalchuk Y, Konnerth A (1996) Long-term potentiation and functional synapse induction in developing hippocampus. Nature 381:71–75.

Eng JK, McCormack AL, Yates JR (1994) An approach to correlate tandem mass spectral data of peptides with amino acid sequences in a protein database. J Am Soc Mass Spectrom 5:976–989.

Ewels P, Magnusson M, Lundin S, Kaller M (2016) MultiQC: summarize analysis results for multiple tools and samples in a single report. Bioinformatics 32:3047–3048.

Geisheker MR et al. (2017) Hotspots of missense mutation identify neurodevelopmental disorder genes and functional domains. Nat Neurosci 20:1043–1051.

Giovannoni F, Quintana FJ (2020) The Role of Astrocytes in CNS Inflammation. Trends Immunol 41:805–819.

Guzman YF, Ramsey K, Stolz JR, Craig DW, Huentelman MJ, Narayanan V, Swanson GT (2017) A gain-of-function mutation in the GRIK2 gene causes neurodevelopmental deficits. Neurol Genet 3:e129.

Hanse E, Seth H, Riebe I (2013) AMPA-silent synapses in brain development and pathology. Nat Rev Neurosci 14:839–850.

Hansen KB, Wollmuth LP, Bowie D, Furukawa H, Menniti FS, Sobolevsky AI, Swanson GT, Swanger SA, Greger IH, Nakagawa T, McBain CJ, Jayaraman V, Low CM, Dell’Acqua ML, Diamond JS, Camp CR, Perszyk RE, Yuan H, Traynelis SF (2021) Structure, Function, and Pharmacology of Glutamate Receptor Ion Channels. Pharmacol Rev 73:298–487.

Harlow EG, Till SM, Russell TA, Wijetunge LS, Kind P, Contractor A (2010) Critical period plasticity is disrupted in the barrel cortex of FMR1 knockout mice. Neuron 65:385–398.

Harris JJ, Jolivet R, Attwell D (2012) Synaptic energy use and supply. Neuron 75:762–777.

Hensch TK (2005) Critical period plasticity in local cortical circuits. Nat Rev Neurosci 6:877-888.

Iossifov I et al. (2014) The contribution of de novo coding mutations to autism spectrum disorder. Nature 515:216–221.

Isaac JT, Nicoll RA, Malenka RC (1995) Evidence for silent synapses: implications for the expression of LTP. Neuron 15:427–434.

Isaac JT, Crair MC, Nicoll RA, Malenka RC (1997) Silent synapses during development of thalamocortical inputs. Neuron 18:269–280.

Ismail V, Zachariassen LG, Godwin A, Sahakian M, Ellard S, Stals KL, Baple E, Brown KT, Foulds N, Wheway G, Parker MO, Lyngby SM, Pedersen MG, Desir J, Bayat A, Musgaard M, Guille M, Kristensen AS, Baralle D (2022) Identification and functional evaluation of GRIA1 missense and truncation variants in individuals with ID: An emerging neurodevelopmental syndrome. Am J Hum Genet 109:1217–1241.

Katz LC, Shatz CJ (1996) Synaptic activity and the construction of cortical circuits. Science 274:1133–1138.

Klein RM, Howe JR (2004) Effects of the lurcher mutation on GluR1 desensitization and activation kinetics. J Neurosci 24:4941–4951.

Kohda K, Wang Y, Yuzaki M (2000) Mutation of a glutamate receptor motif reveals its role in gating and delta2 receptor channel properties. Nat Neurosci 3:315–322.

Li R, Hu K, Liu H, Green MR, Zhu LJ (2020) OneStopRNAseq: A Web Application for Comprehensive and Efficient Analyses of RNA-Seq Data. Genes (Basel) 11.

Li Z, Okamoto K, Hayashi Y, Sheng M (2004) The importance of dendritic mitochondria in the morphogenesis and plasticity of spines and synapses. Cell 119:873–887.

Liao D, Hessler NA, Malinow R (1995) Activation of postsynaptically silent synapses during pairing-induced LTP in CA1 region of hippocampal slice. Nature 375:400–404.

Liao Y, Smyth GK, Shi W (2014) featureCounts: an efficient general purpose program for assigning sequence reads to genomic features. Bioinformatics 30:923–930.

Lin YJ, Huang TN, Hsueh YP (2023) Quantification of the density and morphology of dendritic spines and synaptic protein distribution using Thy1-YFP transgenic mice. STAR Protoc 4:102290.

Liu D, Tao K, Wang Y, Sun Y, Wang S (2026) Mitochondrial-Epigenetic Crosstalk in Autism Spectrum Disorder: Linking Cellular Stress to Synaptic Dysfunction and Treatment Resistance. Molecular neurobiology 63.

McAlister GC, Nusinow DP, Jedrychowski MP, Wuhr M, Huttlin EL, Erickson BK, Rad R, Haas W, Gygi SP (2014) MultiNotch MS3 enables accurate, sensitive, and multiplexed detection of differential expression across cancer cell line proteomes. Anal Chem 86:7150–7158.

Mootha VK et al. (2003) PGC-1alpha-responsive genes involved in oxidative phosphorylation are coordinately downregulated in human diabetes. Nature genetics 34:267–273.

Moya NA, Yun S, Fleps SW, Martin MM, Nadel JA, Beutler LR, Zweifel LS, Parker JG (2023) The effect of selective nigrostriatal dopamine excess on behaviors linked to the cognitive and negative symptoms of schizophrenia. Neuropsychopharmacology 48:690–699.

Myers SJ, Yuan H, Perszyk RE, Zhang J, Kim S, Nocilla KA, Allen JP, Bain JM, Lemke JR, Lal D, Benke TA, Traynelis SF (2023) Classification of missense variants in the N-methyl-d-aspartate receptor GRIN gene family as gain-or loss-of-function. Hum Mol Genet 32:2857–2871.

Nomura T, Taniguchi S, Wang YZ, Yeh NH, Wilen AP, Castillon CCM, Foote KM, Xu J, Armstrong JN, Savas JN, Swanson GT, Contractor A (2023) A Pathogenic Missense Mutation in Kainate Receptors Elevates Dendritic Excitability and Synaptic Integration through Dysregulation of SK Channels. J Neurosci 43:7913–7928.

Pang XP, Hershman JM, Chung M, Pekary AE (1989) Characterization of tumor necrosis factor-alpha receptors in human and rat thyroid cells and regulation of the receptors by thyrotropin. Endocrinology 125:1783–1788.

Patani R, Hardingham GE, Liddelow SA (2023) Functional roles of reactive astrocytes in neuroinflammation and neurodegeneration. Nat Rev Neurol 19:395–409.

Pennington ZT, Dong Z, Feng Y, Vetere LM, Page-Harley L, Shuman T, Cai DJ (2019) ezTrack: An open-source video analysis pipeline for the investigation of animal behavior. Sci Rep 9:19979.

Pickard L, Noel J, Henley JM, Collingridge GL, Molnar E (2000) Developmental changes in synaptic AMPA and NMDA receptor distribution and AMPA receptor subunit composition in living hippocampal neurons. J Neurosci 20:7922–7931.

Rangaraju V, Lauterbach M, Schuman EM (2019) Spatially Stable Mitochondrial Compartments Fuel Local Translation during Plasticity. Cell 176:73–84 e15.

Resnik E, McFarland JM, Sprengel R, Sakmann B, Mehta MR (2012) The effects of GluA1 deletion on the hippocampal population code for position. J Neurosci 32:8952–8968.

Ritchie ME, Phipson B, Wu D, Hu Y, Law CW, Shi W, Smyth GK (2015) limma powers differential expression analyses for RNA-sequencing and microarray studies. Nucleic acids research 43:e47.

Rojas-Charry L, Nardi L, Methner A, Schmeisser MJ (2021) Abnormalities of synaptic mitochondria in autism spectrum disorder and related neurodevelopmental disorders. J Mol Med (Berl) 99:161–178.

Rossignol DA, Frye RE (2012) Mitochondrial dysfunction in autism spectrum disorders: a systematic review and meta-analysis. Mol Psychiatry 17:290–314.

Salpietro V et al. (2019) AMPA receptor GluA2 subunit defects are a cause of neurodevelopmental disorders. Nat Commun 10:3094.

Schwarz LA et al. (2026) Cortical development dynamics across autism spectrum disorder mouse models. Nature.

Shepherd JD, Huganir RL (2007) The cell biology of synaptic plasticity: AMPA receptor trafficking. Annu Rev Cell Dev Biol 23:613–643.

Smyth GK (2004) Linear models and empirical bayes methods for assessing differential expression in microarray experiments. Stat Appl Genet Mol Biol 3:Article3.

Sobolevsky AI, Rosconi MP, Gouaux E (2009) X-ray structure, symmetry and mechanism of an AMPA-subtype glutamate receptor. Nature 462:745–756.

Stevens B, Allen NJ, Vazquez LE, Howell GR, Christopherson KS, Nouri N, Micheva KD, Mehalow AK, Huberman AD, Stafford B, Sher A, Litke AM, Lambris JD, Smith SJ, John SW, Barres BA (2007) The classical complement cascade mediates CNS synapse elimination. Cell 131:1164–1178.

Subramanian A, Tamayo P, Mootha VK, Mukherjee S, Ebert BL, Gillette MA, Paulovich A, Pomeroy SL, Golub TR, Lander ES, Mesirov JP (2005) Gene set enrichment analysis: a knowledge-based approach for interpreting genome-wide expression profiles. Proc Natl Acad Sci U S A 102:15545–15550.

Tabb DL, McDonald WH, Yates JR, 3rd (2002) DTASelect and Contrast: tools for assembling and comparing protein identifications from shotgun proteomics. J Proteome Res 1:21–26.

Ting L, Rad R, Gygi SP, Haas W (2011) MS3 eliminates ratio distortion in isobaric multiplexed quantitative proteomics. Nat Methods 8:937–940.

Turrigiano GG, Nelson SB (2004) Homeostatic plasticity in the developing nervous system. Nat Rev Neurosci 5:97–107.

Tvergaard NK, Tkemaladze T, Stodberg T, Kvarnung M, Tatton-Brown K, Baralle D, Tumer Z, Bayat A (2024) Unraveling GRIA1 neurodevelopmental disorders: Lessons learned from the p.(Ala636Thr) variant. Clin Genet 106:427-436.

Vardalaki D, Chung K, Harnett MT (2022) Filopodia are a structural substrate for silent synapses in adult neocortex. Nature 612:323–327.

Virga DM, Hamilton S, Osei B, Morgan A, Kneis P, Zamponi E, Park NJ, Hewitt VL, Zhang D, Gonzalez KC, Russell FM, Grahame Hardie D, Prudent J, Bloss E, Losonczy A, Polleux F, Lewis TL, Jr. (2024) Activity-dependent compartmentalization of dendritic mitochondria morphology through local regulation of fusion-fission balance in neurons in vivo. Nat Commun 15:2142.

Wang YZ, Perez-Rosello T, Smukowski SN, Surmeier DJ, Savas JN (2024a) Neuron type-specific proteomics reveals distinct Shank3 proteoforms in iSPNs and dSPNs lead to striatal synaptopathy in Shank3B(-/-) mice. Mol Psychiatry 29:2372–2388.

Wang YZ, Castillon CCM, Gebis KK, Bartom ET, d’Azzo A, Contractor A, Savas JN (2024b) Notch receptor-ligand binding facilitates extracellular vesicle-mediated neuron-to-neuron communication. Cell Rep 43:113680.

Webb BT, Trinh H, Breach EA, Foote KM, Binelli E, Swanson GT (2026) Pathological gain-of-function human variants in the GRIK2 kainate receptor gene cause wide-ranging behavioral dysfunction and seizures in mouse models. Neurobiol Dis 218:107226.

XiangWei W et al. (2023) Clinical and functional consequences of GRIA variants in patients with neurological diseases. Cell Mol Life Sci 80:345.

Xu J, DeVries SH, Zhu Y (2020) Quantification of Adeno-Associated Virus with Safe Nucleic Acid Dyes. Hum Gene Ther 31:1086–1099.

Xu J, Contractor A, Zhu Y (2022) Protocol to map multi-transmitter neurons in the mouse brain using intersectional strategy. STAR Protoc 3:101907.

Xu J, Zhu Y, Contractor A, Heinemann SF (2009) mGluR5 has a critical role in inhibitory learning. J Neurosci 29:3676–3684.

Xu J, Zhu Y, Kraniotis S, He Q, Marshall JJ, Nomura T, Stauffer SR, Lindsley CW, Conn PJ, Contractor A (2013) Potentiating mGluR5 function with a positive allosteric modulator enhances adaptive learning. Learn Mem 20:438–445.

Xu T, Park SK, Venable JD, Wohlschlegel JA, Diedrich JK, Cociorva D, Lu B, Liao L, Hewel J, Han X, Wong CCL, Fonslow B, Delahunty C, Gao Y, Shah H, Yates JR, 3rd (2015) ProLuCID: An improved SEQUEST-like algorithm with enhanced sensitivity and specificity. J Proteomics 129:16–24.

